# Using deep generative models for simultaneous representational and predictive modeling of brain and behavior: A graded unsupervised-to-supervised modeling framework

**DOI:** 10.1101/2024.12.23.630166

**Authors:** Kieran McVeigh, Ashutosh Singh, Deniz Erdogmus, Lisa Feldman Barrett, Ajay B. Satpute

## Abstract

This paper uses a generative neural network architecture combining unsupervised (generative) and supervised (discriminative) models with a model comparison strategy to evaluate assumptions about the mappings between brain states and behavior. Most modeling in cognitive neuroscience publications assume a one-to-one brain-behavior relationship that is linear, but never test these assumptions or the consequences of violating them. We systematically varied these assumptions using simulations of four ground-truth brain-behavior mappings that involve progressively more complex relationships, ranging from one-to-one linear mappings to many-to-one nonlinear mappings. We then applied our Variational AutoEncoder-Classifier framework to the simulations to show how it accurately captured diverse brain-behavior mappings,provided evidence regarding which assumptions are supported by the data, and illustrated the problems that arise when assumptions are violated. This integrated approach offers a reliable foundation for cognitive neuroscience to effectively model complex neural and behavioral processes, allowing more justified conclusions about the nature of brain-behavior mappings.

## Introduction

Research in cognitive neuroscience has increasingly used machine learning algorithms and multivariate pattern analysis (MVPA) to model brain-behavior relationships (Haxby, 2012; Norman et al., 2006; Peelen & Downing, 2023; Pereira et al., 2009). These algorithms typically fall into two main types: unsupervised or supervised (Hastie et al., 2001). Unsupervised methods estimate the distributions of brain states derived from measurements during an experiment but without reference to behavior (Fukushima, 1980; Goutte et al., 1999; Han et al., 2019; Huang et al., 2018). Supervised methods learn how brain states predict behavior (e.g. condition labels; Haynes & Rees, 2006; Mitchell et al., 2004; O’Toole et al., 2007; Poldrack et al., 2009; Tong & Pratte, 2012). Discriminative supervised methods, which are the most commonly used supervised methods in cognitive neuroscience, do not estimate the distribution of brain states (generative supervised approaches do, but are limited because they have stringent assumptions about data distributions; Bishop, 2006). In cognitive neuroscience, most studies assume that brain states map to behavior in a linear, one-to-one fashion (Barrett & Satpute, 2013; Poldrack, 2008; Rugg & Thompson-Schill, 2013; Uttal, 2003). If such mapping exists, unsupervised and supervised approaches should lead to converging conclusions (see **Figure 1A**; Azari et al., 2020). Conclusions diverge, however, when the mapping is more complex (**Figure 1B-D**; e.g., Westlin et al., 2023). It could be, for example, that the brain-behavior mapping is not one-to-one, that the brain-behavior mapping is not linearly separable, or that the dimensions of highest variance in the brain data do not relate to behavior. Unfortunately, the ground truth of brain-behavior relationships is rarely, if ever, known in advance, leading to the possibility of incorrect conclusions when scientists use a modeling approach guided by a single set of assumptions that may not be justified (e.g., using a linear discriminative supervised method on brain data that maps to behavior in a nonlinear and/or many-to-one, brain states-to-behavior fashion, see Azari et al., 2020).

**Figure 1.**
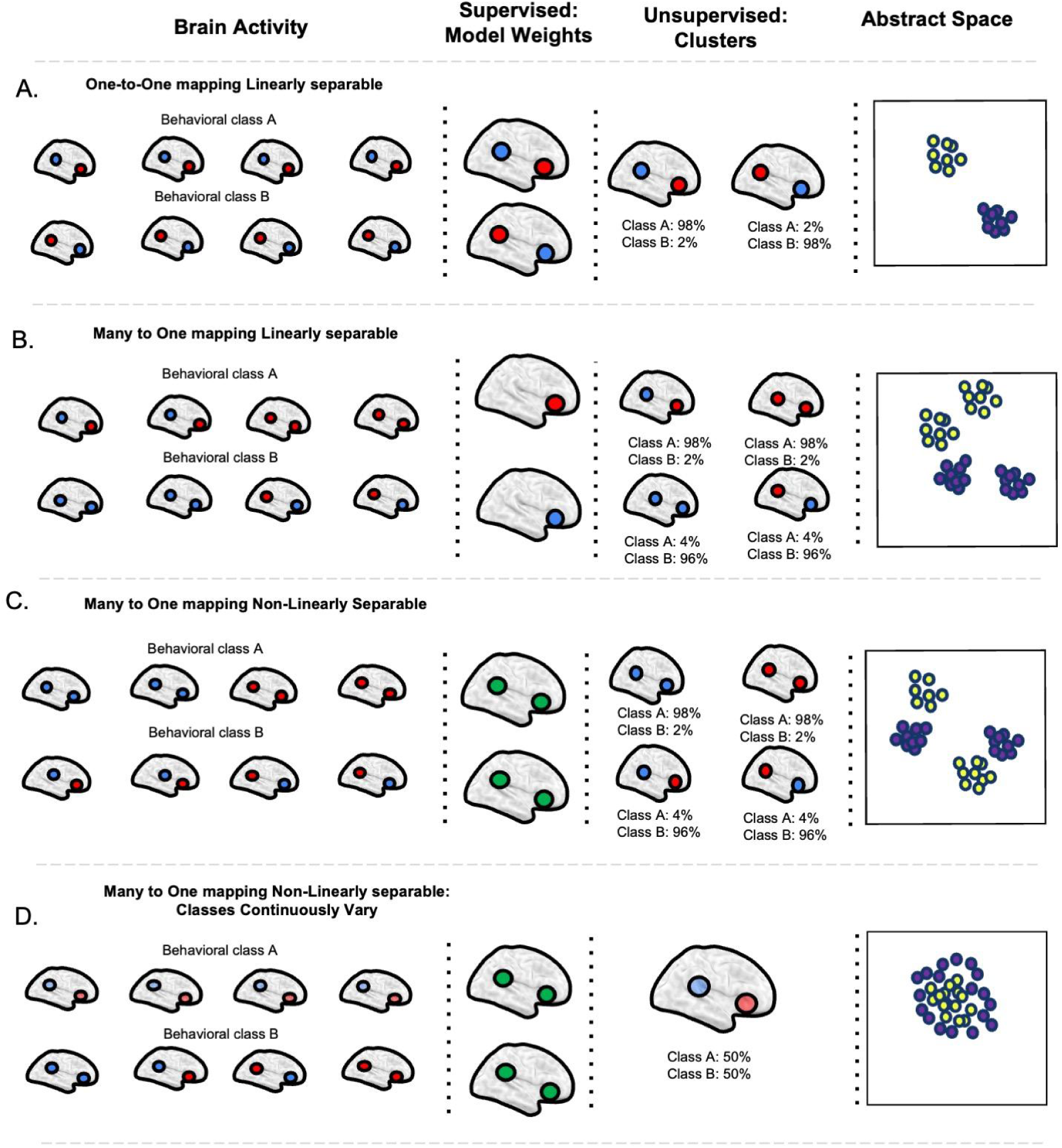
Four hypothetical brain-behavior mappings. This figure provides simplified cartoon examples of possible brain-behavior relationships, and how supervised and unsupervised clustering approaches might model these mappings. **Panel A** shows a simple one-to-one brain-behavior mapping. The left most column depicts brain activity occurring during instances of behavioral class A (e.g. four trials in an fMRI study; top row) and behavior class B (bottom row). A one-to-one mapping assumes that the same pattern of brain activity occurs across instances. The second column represents the hypothetical feature weights of behavioral classes as modeled by a linear discriminative supervised algorithm. For instance the features weights of a logistic regression would show that increases in the anterior region and decreases in the posterior region in activity contribute to a prediction of behavioral class A (and vice versa for behavioral class B). **Cluster finding** These feature weights correspond with the underlying brain states, and the unsupervised clustering findings (shown in the third column). Finally the last column shows the same scenario in a two dimensional brain state space. **Of note under the restrictive assumption of a one-to-one mapping these approaches converge. Panel B** depicts a hypothetical many-to-one brain-behavior mapping, **wherein** there are multiple brain states associated with each behavioral class, but **behavioral classes are still linearly separable**. A linear discriminative supervised algorithm will succeed in classifying behavioral classes from brain data, but it will fail in capturing the multiple brain states associated with the same class (i.e., it will only model regions contributing to classification). In contrast, an unsupervised approach may model the four brain states, albeit does not explicitly relate them to behavior. The final column illustrates this scenario in a brain state space, showing 4 clusters of data and linearly-separable, many-to-one mapping. **Panel C** depicts a many-to-one nonlinearly separable brain-behavior mapping. Here, a linear supervised model would not classify well in this scenario, and so we assume a nonlinear supervised discriminative algorithm for our example. Although a nonlinear model will succeed in classification, they provide little information about the brain states underlying the behavioral classes. We can calculate average importance weights (indicated in green in the third column), which indicate which features are more or less important, but importance weights are not easily relatable to information about brain states (unlike regression weights which, even while not directly modeling brain states, at least provide indications of both sign and magnitude of importance of activity in a brain region). Unsupervised analyses do capture this many-to-one mapping (but again, do not explicitly relate brain states to behavior). The final column illustrates this scenario in a brain state space, showing 4 clusters of data, and a many-to-one nonlinearly separable mapping. **Panel D** depicts another many-to-one, nonlinearly separable brain-behavior mapping (sometimes referred to as the “Anna Karenina” organization; Finn et al., 2020). Instances of behavioral class 1 have a similar pattern of activity whereas instances of class 2 are highly heterogenous. A nonlinear supervised model would successfully classify the behavioral classes, but again the importance weights do not accurately describe the brain states associated with each class. In fact, the average importance weights in this case are identical to those in Panel C even though the brain-behavior mapping is completely different, which further underscores the shortcomings of using feature weights from supervised models to gain insight on brain states. Critically, this scenario also reveals the shortcomings of a fully unsupervised approach. For example, a typical clustering solution would reveal a single cluster, failing to even separate the data into behaviorally relevant classes. The final column illustrates this scenario in a brain state space, showing the Anna Karenina-like organization of data.

In this paper, we introduce a possible solution to this dilemma. We combine unsupervised and discriminative supervised models with a model comparison strategy, and apply this approach on simulated data where the ground truth brain-behavior relationships are known. The combined modeling approach learns a latent space that models the distribution of simulated brain states (similar to unsupervised approaches). At the same time, the latent space contains information that can be used to predict behavior, thereby contributing to brain-behavior characterizations (similar to supervised approaches). We use four simulated datasets that vary in the linearity and homogeneity of the brain-behavior relationship to compare the modeling results for each simulated dataset along a continuum from fully unsupervised to fully supervised. More importantly, we examine what happens to the latent space as a consequence of incorrect assumptions that applied to a dataset during modeling. We further show how our framework can model diverse brain-behavior relationships in a way that fully unsupervised v. fully supervised approaches do not, by comparing the modeling results for each simulated dataset along a continuum from fully unsupervised to fully supervised.

### Supervised Approaches

Supervised algorithms are explicitly designed to estimate a predictive function from brain data to behavioral labels (Bishop, 2006). They have been successfully used to predict labels across multiple psychological domains including memory, social cognition, emotion, language, etc., from whole-brain signal patterns (Pereira et al., 2009; Poldrack et al., 2009; Tong & Pratte, 2012; Weaverdyck et al., 2020). Supervised methods offer two key advantages for making inferences about brain-behavior mappings. First, measurements of brain activity include many sources of variance including noise and supervised approaches use only the fraction of that variance that predicts the behavioral labels, given specific modeling assumptions. Second, brain-behavior relationships may be complex (e.g., nonlinear, many-to-one mappings; **Figure 1C-D**) and supervised approaches can nonetheless learn to predict behavior even in the presence of such complexities (e.g., by using neural network models as a “universal function approximators”; Hornik et al., 1989).

These strengths are directly related to the key weakness of the supervised approach. A complete understanding of brain-behavior relations *requires* modeling the distribution of brain states. Popular supervised algorithms do not explicitly represent the brain data, however. In the absence of such modeling, it is possible to predict behavioral labels with the same solution in datasets that have different brain-behavior relations, obscuring those differences (Clark-Polner et al., 2017; Lindquist et al., 2022). For example, consider **Figure 1A** vs. **Figure 1B**. Both datasets could be modeled with the same discriminative algorithm and distinguish between two classes (open circles above the dark circles), even though the two datasets involve different brain-behavior relations. Discriminative algorithms such as support vector machines or LASSO-PCR model a decision-boundary that is used to distinguish one class from another to predict behavioral labels, called a hyperplane. These hyperplanes are then analyzed to understand brain-behavior relations. Since these hyperplanes fail to model brain states, it is quite possible for these algorithms to estimate the same hyperplane across fundamentally different brain-behavior relations (Kragel et al., 2018; Wang et al., 2024).

As long as the behavioral predictions are successful, it could be argued that understanding the characteristics of the underlying brain behavior relationships (e.g. the distribution of brain states) is unnecessary. This argument comes with some fine print: Because many relationships can be described by the same solution, *inferences* about ground-truth relationships are unjustified. The risk of ill-advised inferences is particularly pronounced in cognitive neuroscience. Researchers often fit a single linear model and, if it performs significantly above chance, interpret it as accurately capturing the brain-behavior relationship. Yet this model assumes—rather than tests—key aspects of the relationship, such as its linearity, possibly obscuring its true nature. Thus, while supervised approaches can be used to develop predictive models, they offer limited insight into the brain-behavior relations that are the main theoretical focus of cognitive neuroscience.

### Unsupervised Approaches

The learning objective of unsupervised algorithms is to model structured variance (i.e., reliable patterns) in the data irrespective of any labels (Hastie et al., 2001). Researchers have used unsupervised models to take an “inside out” approach to brain-behavior mapping wherein models of structured variance in brain data are estimated first, and then the model’s estimates are related to behavioral data such as stimulus features, behavioral responses, etc. (Azari et al., 2020; Buzsáki, 2019; Sennesh et al., 2020; for discussion, see Westlin et al., 2023). Unsupervised approaches offer several strengths over supervised algorithms. First, they explicitly model brain states. That is, they model brain data as distributions (when using clustering algorithms) or as points in a lower dimensional space (when using other dimensionality reduction algorithms). Second, unsupervised approaches allow for more flexible, data-driven conclusions to emerge. Observed clusters or dimensions are not inherently constrained by experimenter-defined condition labels, participant categories, or any other theory-laden assumptions of how brain and behavior “should” relate (e.g., the solution is not constrained by assuming there is a one to one mapping between brain and behavior; Khan et al., 2022; Westlin et al., 2023). The solution is still constrained by theory-laden aspects of study design and sampling, but those issues are better discussed elsewhere (Barrett, 2022; Lee et al., 2021; Dubova & Goldstone, 2023). Here, we focus on analytical choices. For example, unsupervised algorithms could reveal that multiple brain states occur during the trials of a given, experimenter-defined task condition (illustrated in **Figure 1B-D**, Azari et al., 2020; Doyle et al., 2022; Khan et al., 2022; Nakuci et al., 2023), consistent with degeneracy or multiple realizability (Edelman & Gally, 2001; Price & Friston, 2002; Putnam, 1960).

These strengths of unsupervised approach approaches are directly related to a key weakness: They assume that the behavior of interest is related to the dominant dimension of variance in the brain data and therefore strong enough to drive model estimates, but that isn’t always the case (for an illustrative case, see **Figure 1D**). Structured variance in brain data during a task may also include signals that are irrelevant to the behavior of interest (by contrast, supervised approaches only model variance related to behavioral labels). Furthermore, it can be difficult to determine whether the signals driving data-driven brain clusters (or dimensions) are in fact behaviorally-relevant (again, see Azari et al., 2020). Thus, while unsupervised approaches model the brain states themselves, the model estimates are not always useful for understanding the brain-behavior relationship.

### Present Study

In this paper, we use recent advances in neural networks that are designed to simultaneously capture the distribution of brain states (the unsupervised learning objective) and model the relationships of brain states to class labels (the supervised learning objective; see **Figure 2A** for architecture diagram). There are a number of ways to implement these neural networks (e.g., Bzdok et al., 2015; Kingma et al., 2014; Zabihi et al., 2024). We implemented the unsupervised portion of the model using a variational auto-encoder (VAE), which is a neural network that learns a latent (lower-dimensional) representation of fMRI BOLD signal data, similar to principal component analysis (PCA). The VAE consisted of an encoder that compressed the BOLD data into a latent space and a decoder that reconstructed the data from the latent representation (**Figure 2A**). The supervised portion of the model used a classification network, also called a classification head, to predict the behavioral labels from the same latent representation (**Figure 2A**). All components (the encoder and decoder of the autoencoder, and the classification head) were trained jointly in an integrated “VAE-C” model (i.e., variational auto-encoder with a classification head). This joint training optimized the learned latent space to capture the underlying structure of the fMRI data and to enable classification of the data directly from the latent space (see **Figure 2A**). For an example of a VAE-C approach applied to EEG data see Bethge et al., (2022).

**Figure 2.**
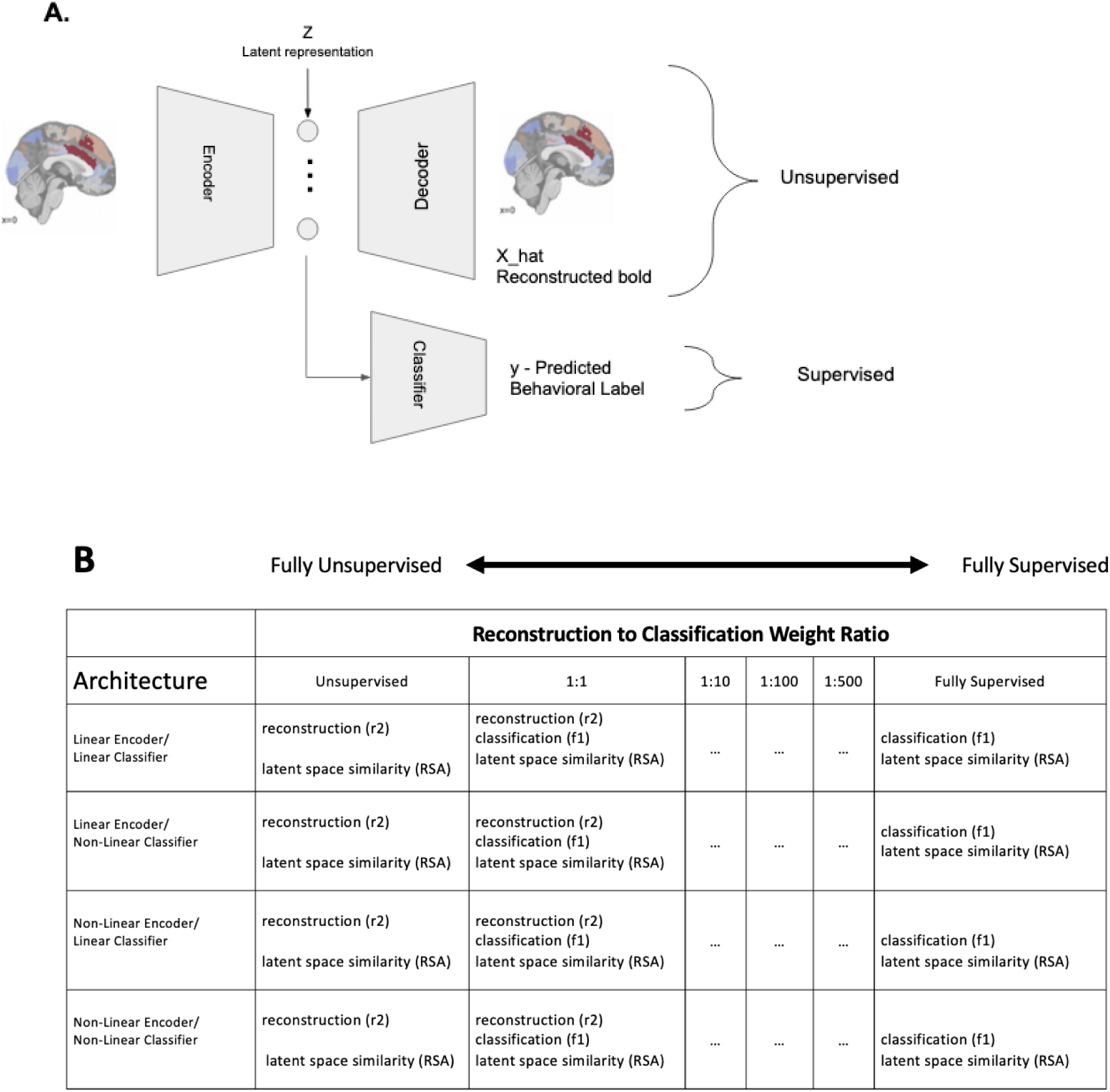
Diagram of network architectures. **A.** shows a schematic of the VAE-C architecture. During training and inference high dimensional fMRI data is input to encoder, and transformed into a lower dimensional latent representation *Z*. This latent representation *Z* is then fed through both decoder and the classifier. The decoder transforms *Z* into X_hat a reconstruction of the inputted bold data while the classifier transforms *Z* into a predicted class label y. The encoding to decoding reconstruction path through the network is an unsupervised analysis that motivates *Z* to model the structure in the inputted bold data. The encoding to classifier path through the network is a supervised analysis that motivates *Z* to contain information relevant to class labels. **B.** A visual representation of our analyses. For each dataset we trained each combination of VAE-C’s with linear or nonlinear encoder and decoders, and linear or nonlinear classifiers - represented as the 2 x 2 table. Additionally we train each combinations of these network architectures at different locations of the supervision continuum, ranging from a fully unsupervised analysis to a fully supervised analysis.

The VAE-C framework allowed us to examine the consequence of introducing increasing supervision during modeling, on a continuum from fully unsupervised to fully supervised (**Figure 2B**). In the fully unsupervised modeling condition, no classification was performed (see left most column of **Figure 2B**). The VAE-C ignored the labels when learning the latent representation of the BOLD data (i.e. there was no weight on the classification objective) and only used the BOLD data itself to estimate the latent space of brain states and reconstruct the data. Both the latent representation and the reconstructed data should recover the ground-truth structure in the simulated BOLD data. At the other end of the continuum in the fully supervised modeling condition (right most column, **Figure 2B**), the behavioral labels strongly shaped the latent representation of the BOLD data (i.e., there was no weight on the reconstruction objective). At various points along the continuum, we successively increased the influence of the labels to examine how the labels impacted the latent representation, the reconstructed data, and the resulting classification: 1 (reconstruction):1 (classification), 1:10, 1:100, and 1:500. (**see Figure 2B**).

In addition, the VAE-C framework contained four different modeling architectures that allowed us to vary whether a network could learn nonlinear relationships to examine the impact on the latent representation (via the encoder function), the reconstructed data (via the decoder function), and the resulting classification (via the classification function). We varied whether the encoder/decoder was restricted to learning a linear relation between the BOLD data and the latent space or whether an encoder/decoder could also learn nonlinear relations, as well as whether the classifier was restricted to learning a linear mapping between the estimated distribution of brain states and the behavioral labels or whether nonlinear mappings could also be estimated. These four architectures were examined for each point along the supervision continuum, resulting in a 2 (linearly or nonlinearly capable auto-encoder) x 2 (a linearly or nonlinearly capable classifier) x W (a weight ratio, W, for the learning objectives from fully unsupervised to fully supervised) framework (see **Figure 2B**).

We then applied this 2×2×W framework to four simulated data sets with different ground truth brain-behavior relationships to examine what happens when modeling assumptions do not respect the ground truth relationships in the modeled data (see Figure 1). We used four ground-truth simulations. Dataset 1 was simulated with a linearly separable, one-to-one brain-behavior mapping, in line with traditional assumptions in cognitive neuroscience (Figure 1A). Dataset 2 was simulated with a linearly separable, but many-to-one brain-behavior relationship (**Figure 1B**). Datasets 3 and 4 embodied the same ground truth relations (nonlinear separable many-to-one brain-behavior mappings) but dataset 3 contained cluster-like mappings whereas dataset 4 contained a continuous mapping (**Figure 1C-D**).

In all simulations, we started with a ground truth latent brain space that contained the distribution of brain states and their groupings (Dataset 1 through 3) or not (Dataset 4). We then generated “observed” BOLD data that was linearly related to that latent space. We also generated “observed” behavioral labels that were either linearly or nonlinearly related to that space. Using the “observed” (i.e., “simulated”) data, we then learned a latent brain space, and from this space reconstructed the BOLD data as well as computed classifications with the labels. This created a unique opportunity to observe the consequences of using different modeling assumptions (embedded in architectures 1 through 4) when brain-behavior mappings are complex (e.g., not one-to-one or nonlinear), and how these consequences might change as the labels had an increasing impact on the learned latent space.

The resulting analysis space was 4 (VAE-C architectures) by 6 weightings (along the supervision continuum from fully unsupervised to fully supervised) by 4 (ground-truth datasets) or 96 analyses. In each analysis, we measured the accuracy of the learned latent space by comparing it to the ground truth latent space for each simulated data set using representational similarity analysis (Kriegeskorte et al., 2008). We measured the accuracy of the reconstructed data by comparing it to the “observed” (simulated) BOLD data using the coefficient of determination (Ezekiel, 1930). We measured the accuracy of the classification by comparing the behavioral labels predicted by the VAE-C analysis to the “observed” (i.e., simulated) behavioral labels using an F1-score (Rijsbergen, 1979).

We predicted and found that, in general, the latent space would be modeled with reasonable accuracy for data that contained straightforward brain-behavior relations (in Dataset 1, the learned and ground-truth latent spaces would converge). We predicted and found that, in general, as the weighting of the classification objective (i.e., the behavioral labels) increased to fully supervised, complex brain-behavior relations (Datasets 2 through 4) would be modeled with distortion or bias (i.e., the learned and ground-truth latent spaces would diverge). As distortions increase, it’s tempting to expect that the reconstructed BOLD data would correspondingly diverge from the “observed” (simulated) BOLD data, but to some extent, this is not what we observed. Instead, as we show, the data reconstructions were largely robust to violated modeling assumptions because more than one latent space produced the same reconstructed data. Also, all functions (encoding, decoding, and classification) proceeded simultaneously and were jointly optimized, such that the decoding function accommodated the distorted space when reconstructing the BOLD data. We predicted and found that all models classified well only when their linearity assumptions matched the ground truth in the simulated data, with one exception as we explain below. Taken together, these analyses make clear that neither supervised or unsupervised modeling alone is sufficient and modeling assumptions should always be tested rather than overlooked.

## Methods

### Modeling Framework

The VAE-C framework learns latent representations which contain information about both the original fMRI signal and the behavioral class associated with the observation. Below we formally describe the more general modeling framework and how our VAE-C architecture implements this framework. We begin by defining our set of variables. Let *D*={*X,Y*}_n_ be the input dataset where *X ∈ R^d^* which represents the matrix of fmri data and *Y ∈ R* represents the behavior label for *n* observations. Let *z ∈ R_d_* be the random variable that describes the latent distribution. We assume that the latent distribution of the generative model is Gaussian {0,*I*}. We specified the learning objective to be,

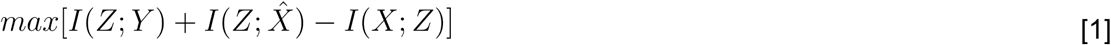

This equation describes a constrained optimization that makes it possible for us to learn a latent representation *Z* such that it contains information about both the class labels *Y* and the neural data *X*. More formally, maximizing the mutual information, *I*, between *Z* and *Y* ensures the latent representation *Z* contains information about the class labels. Maximizing the mutual information between *Z* and [inline1] (i.e. the reconstruction of *X* from *Z*, wherein the optimal case [inline2] = *X*), optimizes the latent representation to be able to reconstruct the data accurately ensuring *Z* contains information about the *X* (the fMRI data). If we only included these first two terms in our model, the model is at greater risk of overfitting the data and generating an uninterpretable latent space. Hence, the final term minimizes mutual information between *Z* and *X*, which regularizes the learned *Z*, taking inspiration from the information bottleneck theory (Tishby et al. 2000).

### VAE Implementation

To implement this broader modeling framework, we used a variational autoencoder with a classification head neural network (VAE-C). The variational autoencoder portion of this network consists of an encoder, and a decoder, implements a generative model of the brain data ( *X*) accomplishing the unsupervised learning objective. While the classifier network from the VAE’s latent space (See **Figure 2A**) accomplishes the supervised learning objective. We use the following notation for describing the parameters of the networks [θ,ϕ,γ] respectively.

Theta is the set of parameters for the encoder, phi is the set of parameters for the decoder, and gamma is the set of parameters for the classifier. We use a VAE derived loss for learning the parameters for [θ,ϕ] which consist of reconstruction loss and a KL (Kulback-leibler) divergence term. Simultaneously we also minimize the cross entropy loss which serves as the proxy for maximizing *I*(*Z;Y*). The objective in [1] can be expressed as the following loss to learn the model parameter set [θ,ϕ,γ],

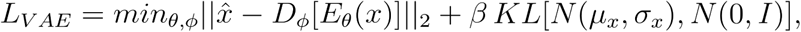

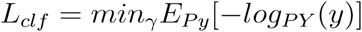

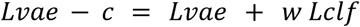

### Network Architecture

To implement our framework, we first specified four VAE-C architectures that varied in the linearity or nonlinearity of the autoencoder and classifier components (**Figure 2B**), that is: (i) linear-autoencoder and linear-classifier, (ii) linear-autoencoder and nonlinear-classifier, (iii) nonlinear-autoencoder and linear-classifier, (iv) nonlinear-autoencoder and nonlinear-classifier. The linear-autoencoders consisted of two fully connected linear layers, one transforming the input ( *X*) to the latent representation ( *Z*), and the other transforming the latent representation ( *Z*) to the reconstruction of the input (X_hat). This architecture is only capable of learning linear encoding and decoding functions. The nonlinear autoencoder adds an additional fully connected hidden layer (size 25), with a tanh activation function, to both the decoder and encoder. The linear-classifier consisted of a single, fully connected linear layer between the latent space ( *Z*) and the class labels ( *Y*). This architecture is only capable of learning a linear function to map from the latent space to class labels. For the nonlinear classifiers, we used two hidden layers between the latent space and the class labels with ReLU activation functions. Additional details and justification for the hidden layer sizes and architecture are located in the hyperparameters section.

### Weighting Learning Objectives

The above equations formally describe how our framework integrates reconstruction and classification objectives. This integration enables a key feature of our architecture - a classification weight - to examine how the modeling framework performs when emphasizing one or the other objective. At one end of the continuum, a classification weight of zero changes the network to be fully unsupervised, which allows us to examine the distribution of the learned *Z* when only fMRI measurements are taken into account. At the other end of the continuum, a reconstruction weight of zero changes the network to be fully supervised, which reveals the brain representations the network learns when functioning as a supervised classifier.

Of particular theoretical interest we can examine how the latent space of the network changes as we gradually adjust the weights of the network to move from a fully unsupervised analysis to a fully supervised analysis. While in relatively simple scenarios we expect the networks learned representations to be consistent across a variety of weights, of greater interest is how in more complex brain behavior mapping (i.e. many-to-one and nonlinear) is understanding how a networks architecture and weights on learning objective interact to shape the learned latent space.

### Simulated Data

We aimed to test our model in four different data simulations that span hypothetical mappings as shown in Figure 1. Dataset 1 was simulated using the conventional assumption of one (brain state) to one (behavior) mapping. Instances (i.e, trials) from behavioral class “A” were sampled from one brain state distribution, whereas those from behavioral class “B” were sampled from another brain state distribution (**Figure 1A**). Latent brain states, which are represented as points in a 2-dimensional space, were sampled from two distinct Gaussian distributions. For each of these gaussians we sampled 500 data points. We then generated behavioral class labels and fMRI data from each point. Behavioral labels were generated by assigning all data points from each gaussian to a given class label (i.e., “A” or “B”). fMRI data from these latent data points was generated by projecting observations from the 2d latent space to the 3d fMRI space (represented as 100 parcels, or regions of interest) using a randomly generated linear function.

To generate the other simulated datasets, this same general procedure was modified as follows. Dataset 2 was simulated using the assumption of a many (brain state) to one (behavior)linearly-separable mapping. Two different brain state distributions were used to generate instances for class “A”, and two different brain state distributions were used to generate instances for class “B” (**Figure 1B**). Latent brain states were sampled from four distinct Gaussian distributions. The simulation generated fMRI and behavioral data that are consistent with a linearly separable, but many-to-one, brain-behavior relationship. Dataset 3 was simulated using the assumption of a many-to-one nonlinearly-separable, brain-behavior mapping with a cluster-like structure (**Figure 1C**). Data was sampled from four gaussians distributions, however the fMRI and behavioral data were simulated such that there was a nonlinearly separable many-to-one, brain-behavior relationship. Dataset 4 was another version of the many-to-one nonlinearly-separable, brain-behavior mapping but with a continuous or dimensional structure (**Figure 1D**). We generated a single distribution of brain states and assigned the 50% of the instances with the greatest similarity to the mean to behavioral class A, and all other instances to behavior class B. For additional details on sampling from the data generating distribution for each simulation, see supplemental materials. Here it’s worth noting that since we did not vary assumptions about the latent space is projected to fMRI observations (i.e. we used the same linear function to project the latent simulated data to the fMRI space) we expect all models to be able to reconstruct the fMRI data nearly perfect at baseline.

### Network Training and Evaluation

All networks were trained with the Adam optimization technique starting with an initial learning rate of.0001. Networks were trained for 1500 epochs. Data was split 80/20 into train and test. To account for the auto-encoder style infrastructure of the networks, where the smallest dimensional layer of the network occurred in the middle of the network, the first 50 epochs of all networks were trained with only the reconstruction and kl divergence objective functions, after which networks were trained with the reported weights on the classification and reconstruction objectives.

For each network, we evaluated reconstruction performance using the coefficient of determination (R²), the classification performance using the F-score, and the representational similarity analysis (RSA) to compare the structure of the learned latent space with the ground truth. We used the F1-score over accuracy as the F1-score balances precision (the proportion of true positive predictions among all positive predictions) and recall (the proportion of true positive predictions among all actual positives). We used RSA because of its common application in cognitive neuroscience and the low dimensionality of our latent spaces also allowed for qualitative comparisons via visual inspections. When calculating the similarity matrices for our RSA analyses we used the euclidean distance as our distance metric and compared the similarity of the matrices with the Pearson’s correlation coefficient. Network performance metrics were calculated at the end of the 1500 epochs on the left out testing data.

## Results

### Overview of Analyses

A summary of our analyses is presented in **Table 1**. The correspondence or “match” of the ground truth brain-behavior relationships in simulated data of and the assumptions built into the architectures in the framework is shown in each cell in the table. As seen in **table 1**, the main mismatches between the ground-truth and VAE-C architectures occur in terms of whether the classification head assumes linear or nonlinear relationships between the brain states and the labels. This distinction corresponds to the type of brain-behavior mapping simulated (linearly or nonlinearly separable). The linearity or nonlinearity of the auto-encoder is of less importance to the current analyses as all relations between the simulated latent and “observed” fMRI data are linear.

**Table 1.**
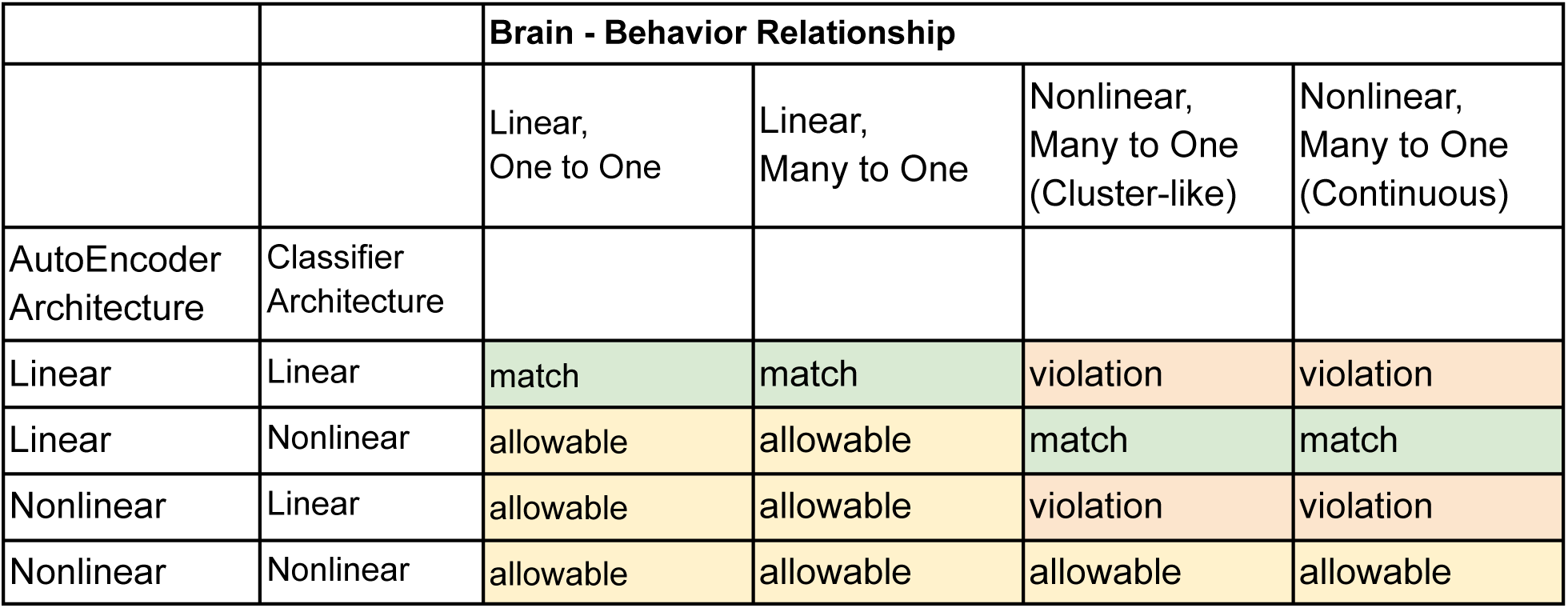
Modeling Assumptions and Simulated Brain-Behavior Relationships. Modeling architectures (along rows) had either linearly or nonlinearly capable encoders and/or classifiers. The brain-behavior relationships in the simulated datasets (along columns) were linearly or nonlinearly separable and also had one-to-one or different versions of many-to-one mappings. Model architectures can “match” (green highlighted cells) or “violate” (red highlighted cells) the data generative process of the ground truth. “Allowable” (yellow highlighted cells) architectures are flexible enough to model the data even if they are not a direct match in terms of the linearity of the encoder and/or the classifier to the data generative process. Whether an architecture is a “match” for a brain-behavior relationship depends on two factors. The first factor is the assumptions of the classifier. Linearly capable classifiers match linear, but violate nonlinear, brain-behavior relationships whereas nonlinearly capable classifiers match nonlinear relationships. The second factor is the assumptions of the encoder. For simplicity, we simulated the “observed” BOLD data using a linear encoding function from the latent space to the brain space. This was helpful for limiting our study to four ground-truth datasets and to maintain our focus on complex brain-*behavior* relationships. But as a consequence, architectures with a nonlinear encoder - regardless of whether they are paired with linear or nonlinear classifiers - can only be allowable (or a violation) and not a match. Intriguingly, the model architectures do not have straightforward matches or mismatches for one-to-one or many-to-one brain behavior mappings. That aspect of brain-behavior mappings is informed in our VAE-C by examining how the latent space is shaped with greater weight on supervision.

### Dataset 1: Linearly Separable, One-to-One Mapping

Dataset 1 was simulated to represent linearly separable Gaussian distributions, such that each behavioral label mapped to one and only one grouping of brain states. This is the brain-behavior mappings most commonly assumed in the majority of analytical approaches used in cognitive neuroscience. The linear encoder/linear classifier architecture contained assumptions that best matched the ground truth represented in Dataset 1 (**Table 1**, column 1). As expected, the learned and ground truth latent state estimates generally converged (i.e. brain states associated with each behavioral label were represented as single grouping in the latent spaces). As the weighting of the labels increased along the supervision continuum, the learned latent space was distorted along the axis perpendicular to the classification boundary (see **Figure 3**), as reflected in decreased RSA scores for the more heavily supervised models (**Table 2**). This distortion did not affect the accuracy of the data reconstruction or classification, as indexed by the R2 and the F1-score values, which remained high across the supervision continuum (R^2^s ∼ 1.00, F1-Scores ∼.95 **Table 2**). This same pattern of results was present for all VAE-C architectures (**Table 2**), as expected, because none of the other VAE-C architectures contained assumptions that violated the linearly separable, one-to-one mapping in Dataset 1.

**Figure 3.**
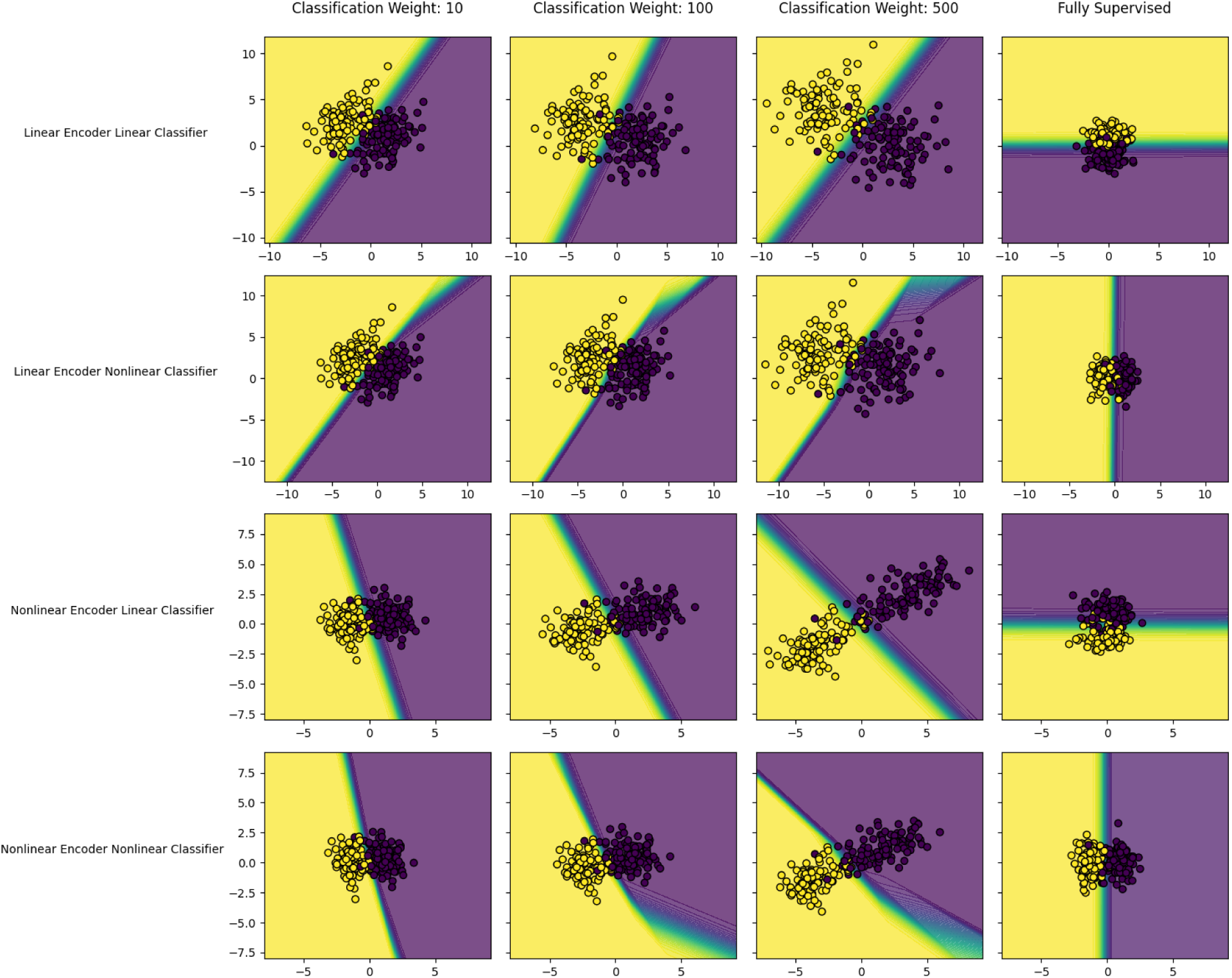
Learned Latent Spaces for Dataset 1. Each panel is the learned latent space for a different iteration of the VAE-C model. Rows correspond to different architectures, while columns correspond to locations on the supervision continuum, with classification weights from left to right (10, 100, 500, and fully supervised). Each dot is the learned latent representation of a simulated fMRI observation. The color of the dot is the “observed” (simulated) behavioral label for that fMRI observation. The background color of each panel represents the models’ prediction of behavior labels for that location in the latent space. Model predictions are probabilistic so areas in which one class is not highly probable show graded coloration. In dataset 1 shown here there is relatively little difference in classification performance and latent representations learned across architectures and classification weights, although slight distortions are seen as the degree of supervision increases.

**Table 2.** Columns of this table are the location on the unsupervised to the supervised continuum models were trained at. This starts with fully unsupervised models in the left columns and increases to fully supervised in the rightmost columns, in between the ratio of the unsupervised to supervised weight is listed. Rows are grouped by the VAE-C architecture a model was trained with. Within each architectural group rows correspond to models performance in terms of reconstruction (R Squared), classification (F1-Score), and similarity of the learned and ground truth latent space (Representational Similarity Analysis; RSA). RSA scores are reported as the Pearson correlation of the ground truth and learned latent space for each model. In this dataset all models perform well in terms of reconstruction, and classification. RSA is high for all unsupervised and semi-supervised models, and shows a moderate decrease in fully supervised models across all architectures.

| **Dataset 1. One-to-one, linearly separable brain-behavior relationship.** |  |  |  |  |  |  |
| --- | --- | --- | --- | --- | --- | --- |
| **Classifier Weight** | **Fully Unsupervised** | **1** | **10** | **100** | **500** | **Fully Supervised** |
| ***Linear Autoencoder, Linear Classifier*** |  |  |  |  |  |  |
| R Squared | 1.00 | 1.00 | 1.00 | 1.00 | 1.00 |  |
| F1-Score |  | 0.96 | 0.95 | 0.96 | 0.96 | 0.91 |
| Accuracy |  | 0.96 | 0.95 | 0.96 | 0.96 | 0.92 |
| RSA | 0.98 | 0.98 | 0.99 | 0.99 | 0.98 | 0.55 |
| kl-divergence | 0.06 | 0.06 | 0.06 | 0.07 | 0.10 | 0.01 |
| ***Linear Autoencoder, Nonlinear Classifier*** |  |  |  |  |  |  |
| R Squared | 1.00 | 1.00 | 1.00 | 1.00 | 1.00 |  |
| F1-Score |  | 0.96 | 0.96 | 0.95 | 0.95 | 0.94 |
| Accuracy |  | 0.96 | 0.96 | 0.95 | 0.95 | 0.94 |
| RSA | 0.98 | 0.98 | 0.98 | 0.99 | 0.99 | 0.49 |
| kl-divergence | 0.06 | 0.06 | 0.06 | 0.06 | 0.10 | 0.01 |
| ***Nonlinear Autoencoder, Linear Classifier*** |  |  |  |  |  |  |
| R Squared | 1.00 | 1.00 | 1.00 | 1.00 | 0.99 |  |
| F1-Score |  | 0.95 | 0.94 | 0.95 | 0.94 | 0.91 |
| Accuracy |  | 0.95 | 0.94 | 0.95 | 0.95 | 0.92 |
| RSA | 0.99 | 0.99 | 0.97 | 0.92 | 0.87 | 0.52 |
| kl-divergence | 0.04 | 0.04 | 0.04 | 0.05 | 0.09 | 0.01 |
| ***Nonlinear Autoencoder, Nonlinear Classifier*** |  |  |  |  |  |  |
| R Squared | 1.00 | 1.00 | 1.00 | 1.00 | 0.99 |  |
| F1-Score |  | 0.94 | 0.95 | 0.95 | 0.95 | 0.92 |
| Accuracy |  | 0.95 | 0.95 | 0.95 | 0.95 | 0.92 |
| RSA | 0.99 | 0.99 | 0.99 | 0.96 | 0.89 | 0.50 |
| kl-divergence | 0.04 | 0.04 | 0.04 | 0.04 | 0.06 | 0 |

### Dataset 2: Linearly Separable, Many-to-One Mapping

Dataset 2 was simulated to represent linearly separable Gaussian distributions, like Dataset 1, except that each behavioral label mapped to more than one distribution of brain states. Similar to Dataset 1, the linear encoder/linear classifier architecture contained assumptions that best matched the ground truth represented in Dataset 2, although none of the other VAE-C architectures contained assumptions that violated that ground truth (**Figure 1**, column 2).

As expected, the fully unsupervised model learned a latent space that converged with the ground truth space (Pearson’s r =.95). The learned latent state became increasingly distorted along the axis perpendicular to the classification boundary as the semi-supervised modeling moved towards fully supervised, as observed for Dataset 1 (**see Figure 4**). The fully supervised model produced the most distortion (Pearson’s r =.24); instead of modeling two distinct groupings for each behavioral label, the fully supervised variant of the the linear encoder/linear classifier architecture produced one grouping for each behavioral label, essentially producing a linearly separable, one-to-one mapping (see **Figure 4** last column). Despite these distortions, data reconstruction and classification remained accurate (**Table 2**). This same pattern of results was present for all VAE-C architectures (**Table 2**), as expected.

**Figure 4.**
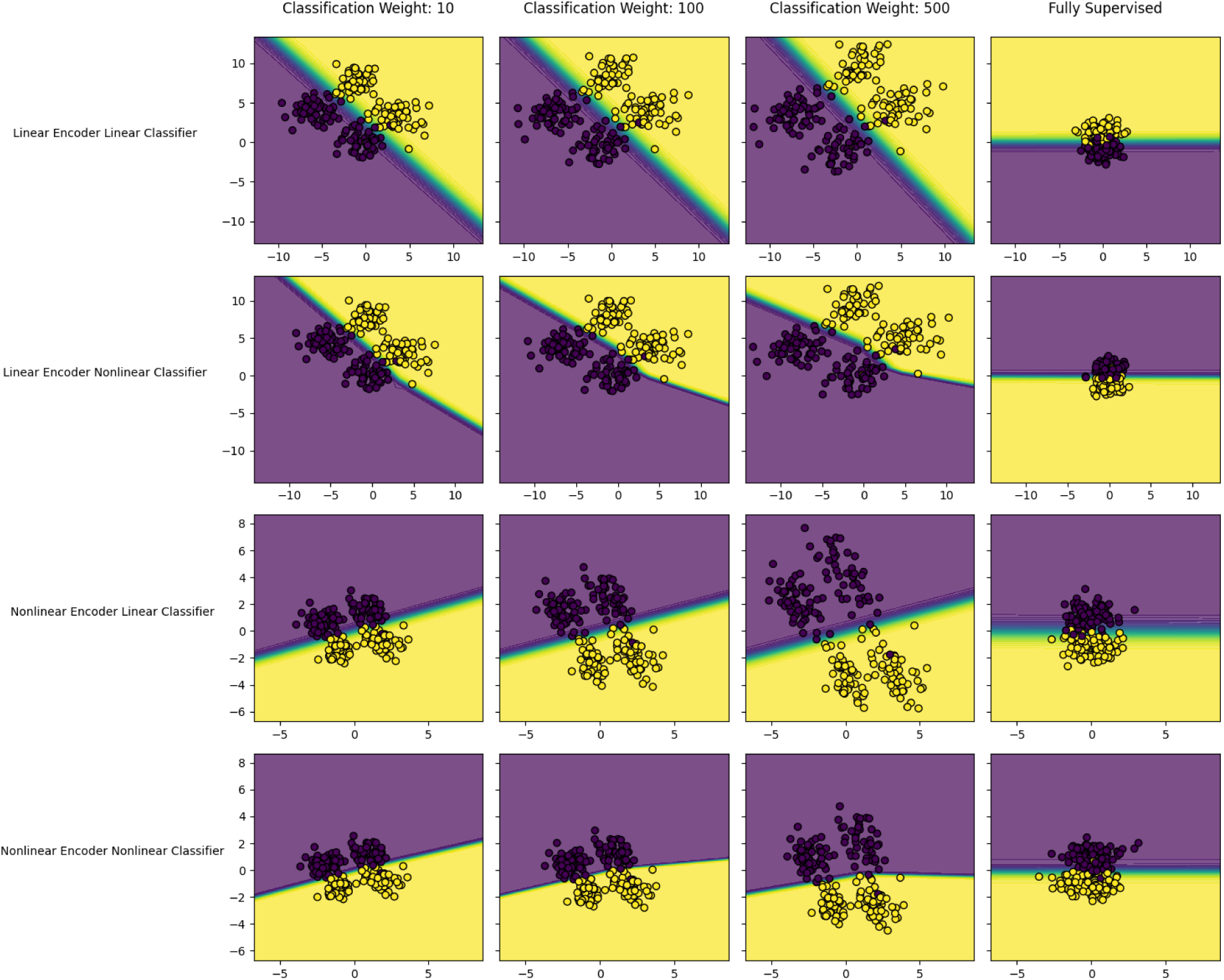
Learned Latent Spaces for Dataset 2. Each panel is the learned latent space for a different iteration of VAE-C model. Rows correspond to different architectures, while columns correspond to locations on the supervision continuum, with classification weights from left to right (10, 100, 500, and fully supervised). Each dot is the learned latent representation of a simulated fMRI observation. The color of the dot is the “observed” (simulated) behavioral label for that fMRI observation. The background color of each panel represents the models’ prediction of behavior labels for that location in the latent space. Model predictions are probabilistic so areas in which one class is not highly probable show graded coloration. In Dataset 2 the main change to the latent space is seen between columns 3 and 4 as models transition to fully supervised models. fully supervised models fail to represent the many to one mappings present in the ground-truth simulated data, despite accurately classifying the brain states.

### Dataset 3: Nonlinearly Separable, Many-to-One Mapping (Discrete)

The linear encoding/nonlinear classifier architecture best matched the ground truth represented in Dataset 3 (**Figure 2**, column 3). Dataset 3 was simulated to represent a many-to-one mapping, such that each behavioral label mapped to more than one distribution of brain states, except that the mapping was nonlinear. As expected, the fully unsupervised model learned a latent space that converged with the ground truth space (Pearson’s r =.98). The semi-supervised models performed similarly. The fully supervised model produced the most distortion (Pearson’s r =.41) in the learned latent space; instead of modeling two distinct distributions for each behavioral label, the fully supervised variant produced one distribution for each behavioral label, essentially creating a linearly separable, one-to-one mapping from the latent space, instead of a nonlinearly separable, many to one mapping (see **figure 5** last column). The classification accuracy for the fully supervised model was similarly low (F1-Score =.66). Data reconstruction accuracy was not computed for the fully supervised model (for which data reconstruction was irrelevant), but was high for all other variants of this architecture (**Table 4**). All semi-supervised variants classified well (F1-Score =.95), with the 1:1 variant being somewhat less accurate (F1-Score =.75).

**Figure 5.**
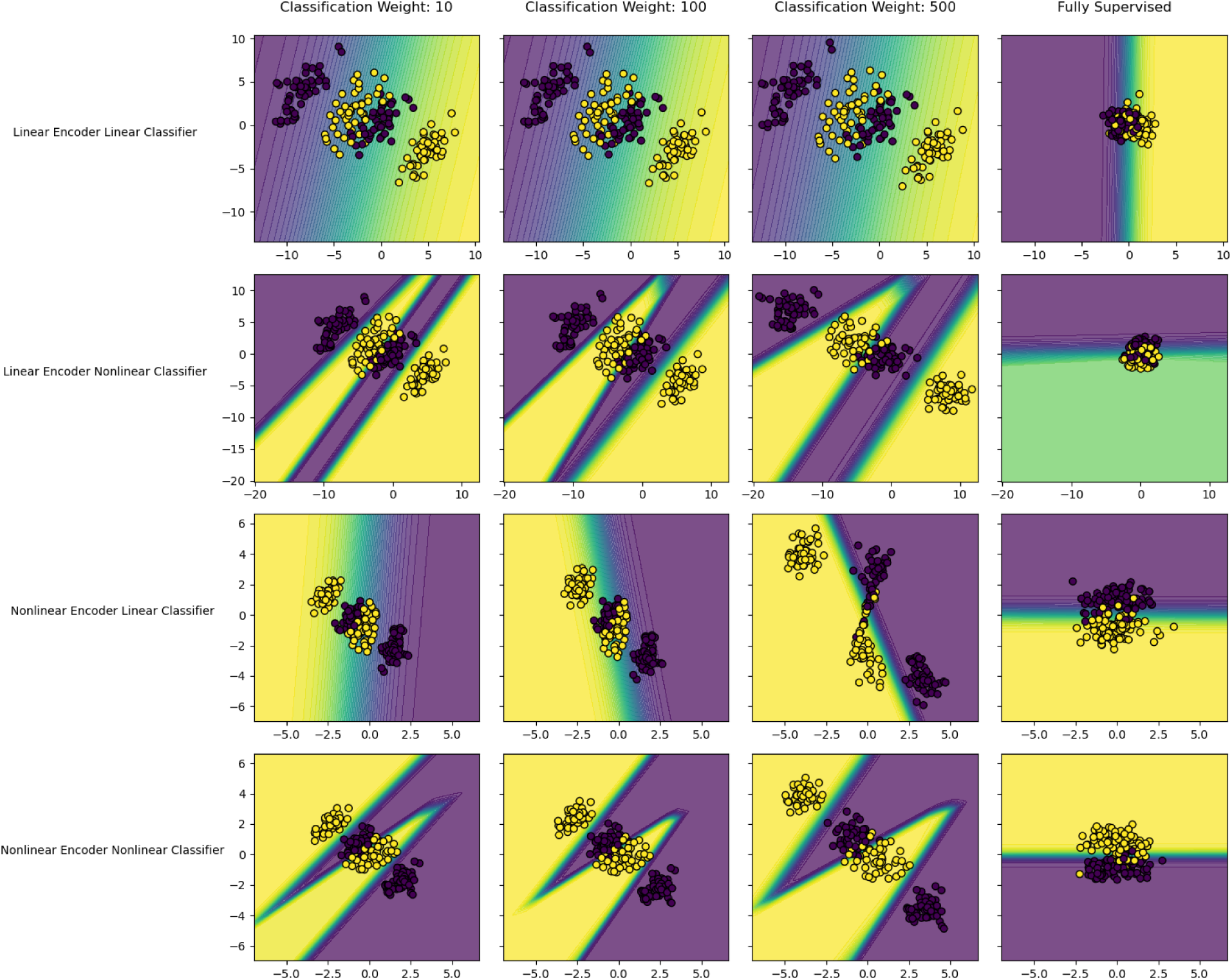
Learned Latent Spaces for Dataset 3. Each panel is the learned latent space for a different iteration of VAE-C model. Rows correspond to different architectures, while columns correspond to locations on the supervision continuum, with classification weights from left to right (10, 100, 500, and fully supervised). Each dot is the learned latent representation of a simulated fMRI observation. The color of the dot is the “observed” (simulated) behavioral label for that fMRI observation. The background color of each panel represents the models’ prediction of behavior labels for that location in the latent space. Model predictions are probabilistic so areas in which one class is not highly probable show graded coloration. For dataset 3, the latent spaces change between columns 3 and 4 as models transition to fully supervised models. Fully supervised models latent spaces fail to represent the many to one mappings present in the ground-truth simulated data. Next for the linear encoder, linear classifier (first row) the graded coloration and poor classification performance demonstrates the model’s inability to learn the nonlinear separation between behavioral labels. Finally for the nonlinear encoder, linear classifier, the latent space becomes progressively distorted along the supervision continuum as the nonlinear encoder learns a nonlinear encoding function to introduce linear separability between brain states associated with different behavioral labels.

**Table 3.**
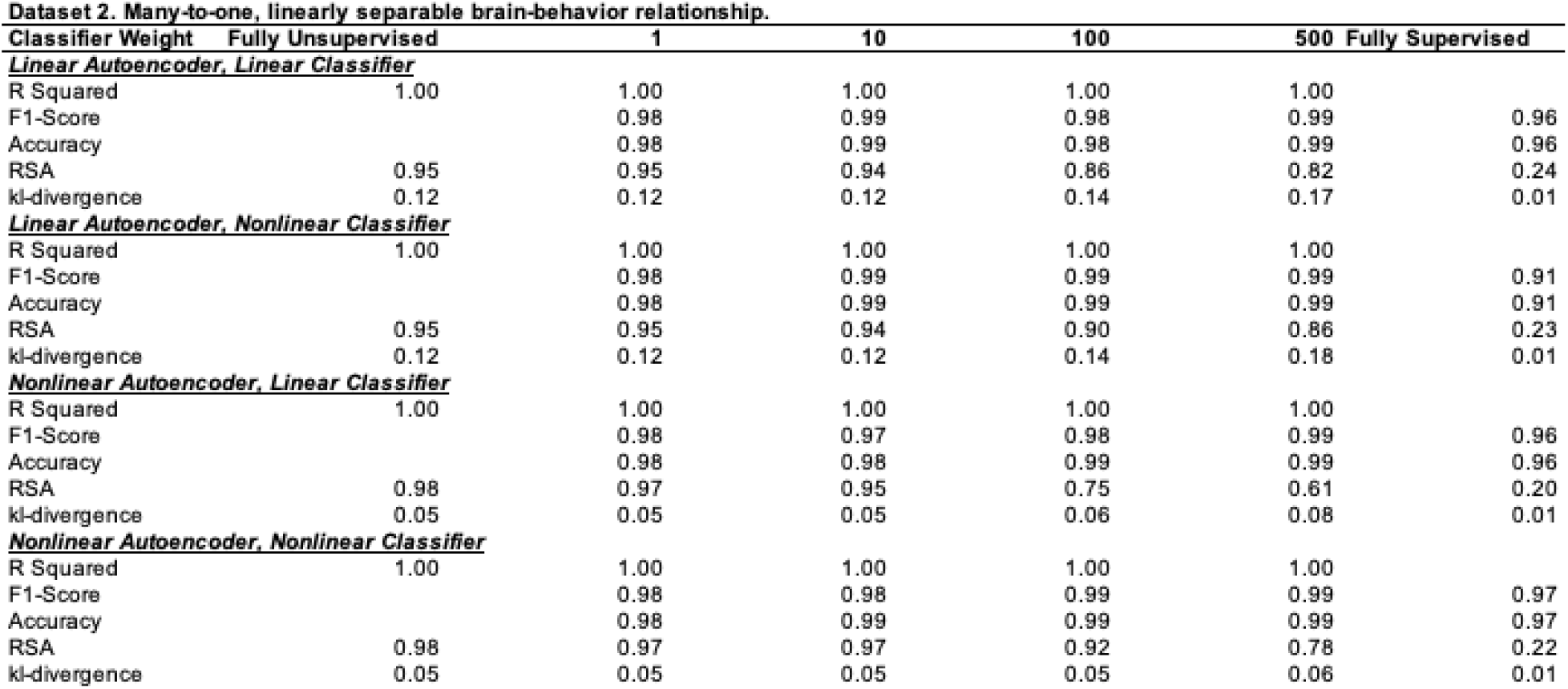
Columns of this table are the location on the unsupervised to the supervised continuum models were trained at. Rows are grouped by the VAE-C architecture a model was trained with. Within each architectural group rows correspond to models performance in terms of reconstruction (R Squared), classification (F1-Score), and similarity of the learned and ground truth latent space (Representational Similarity Analysis; RSA). For dataset 2 all models reconstructed and classified the data accurately. As the weight on the supervised training objective increased to fully supervised the representational similarity of models learned latent representations to the ground truth showed large decrements. Suggesting in the fully supervised case models failed to capture the many to one brain-behavior mappings in dataset 2. Interestingly for many of the semi-supervised models further towards the supervised end of the continuum which showed some distortions to the latent space (RSA scores), the reconstruction was unaffected.

**Table 4.**
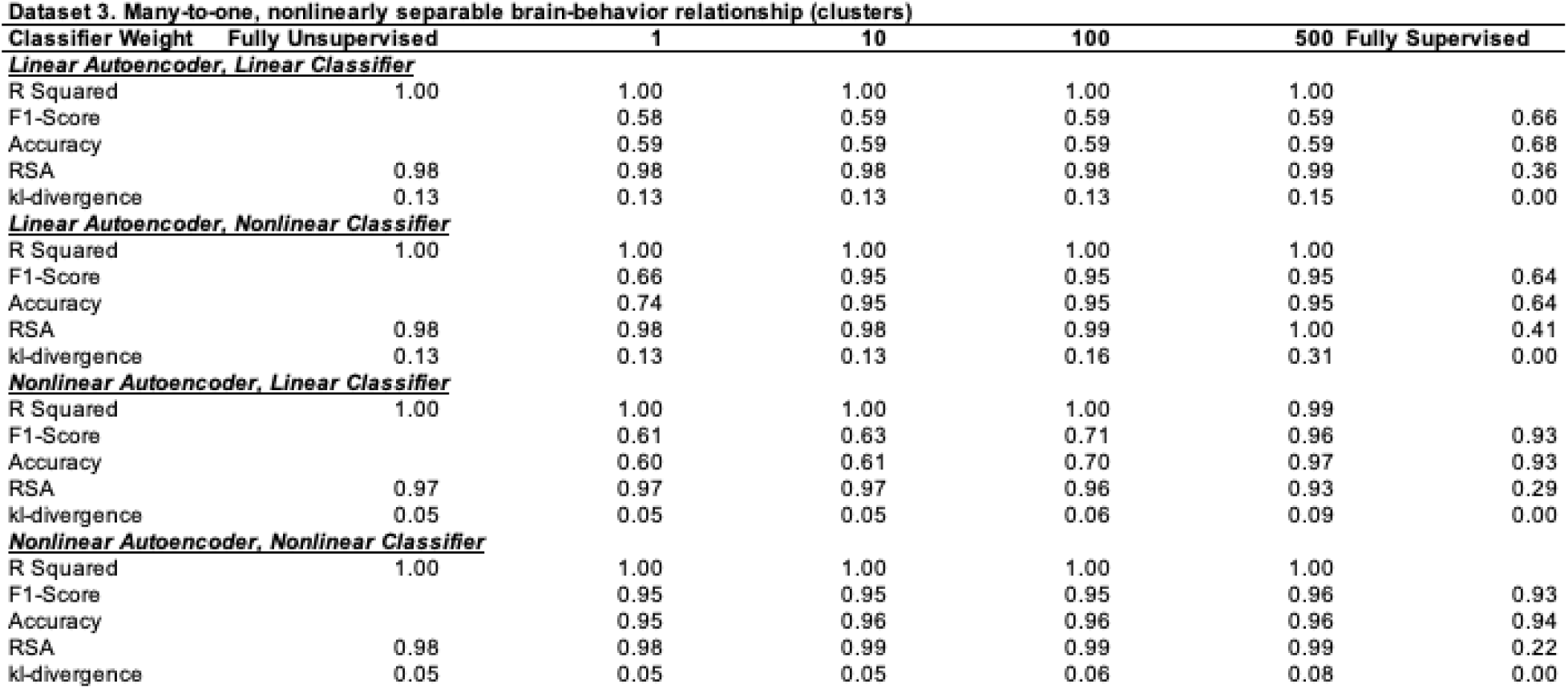
Columns of this table are the location on the unsupervised to the supervised continuum models were trained at. Rows are grouped by the VAE-C architecture a model was trained with. Within each architectural group rows correspond to models performance in terms of reconstruction (R Squared), classification (F1-Score), and similarity of the learned and ground truth latent space (Representational Similarity Analysis; RSA). In dataset 3 all models continued to reconstruct well. Models with nonlinear classifiers generally classified well. All unsupervised and semi-supervised models performed well in terms of Representational Similarity, but fully supervised models performed extremely poorly. The nonlinear autoencoder, linear classifier model showed an interesting set of results, as the classification weight increased the F1-Score increased but representational similarity decreased, suggesting with increasing classification weight this model learned distorted latent spaces to accommodate the linear classifier.

The same pattern of results was generally observed for the architecture that was next best in terms of matching assumptions for Dataset 3: the nonlinear encoding/nonlinear classifier architecture (see Figure 2, column 3). The classifier assumption matched the ground-truth relation between the ground-truth latent space and the “observed” behavioral labels, whereas the nonlinear encoder assumption was capable (but not optimized) for learning the linear encoding relationship between ground-truth latent space and the “observed” (i.e., simulated) BOLD data (**Table 4**). The one exception was the classification accuracy for the fully supervised model, which was high (F1-Score =.93).

The two remaining VAE-C architectures (with linear classifiers) explicitly violated the ground-truth, nonlinear relations between the latent space and the behavioral labels in Dataset 3. The fully unsupervised versions of these two architectures produced accurate learned latent spaces. The fully supervised versions of these two architectures produced distortions in the learned latent space (**Table 4**), but this distortion was unlikely the result of assumption violations because they also emerged in the prior two architectures (which did not explicitly violate any assumptions). The remaining pattern of results for the semi-supervised version of these architectures was complex.

The semi-supervised variants of VAE-C architecture with a linear encoder and a linear classifier learned accurate latent spaces and accurately reconstructed the BOLD data, but their classification accuracy was reduced (F1-Score =.60). The semi-supervised version of the nonlinear encoder combined with a linear classifier learned accurate latent spaces at lower degrees of supervision, but their classification accuracy was reduced. At higher weightings, they produced slight inaccuracies in the learned latent space but, paradoxically, produced high classification accuracy **Table 4**). This effect was a result of this specific architecture (nonlinear encoder/linear classifier). To achieve high classification this architecture requires a latent space where the classes are linearly separable. To achieve this linear separability in dataset 3 (nonlinearly separable many to one mapping) required the nonlinear encoder to learn a distorted latent space (see **figure 5** third row). Despite this distortion to the latent space the reconstruction performance was essentially unaffected.

### Dataset 4: Nonlinearly Separable, Many-to-One Mapping (Continuous)

The results for Dataset 4 largely mirrored those for Dataset 3. Increasing weight on the classification objective increasingly shaped the learned latent space, that space becomes distorted but with little decrement for classification accuracy (and, as we observed in Dataset 3, classification accuracy can even improve, see **Table 5**, **Figure 6**). Distortions in the latent space are associated with slightly less accurate data reconstructions (on the order of a few percentages of variance explained see **Table 5**).

**Figure 6.**
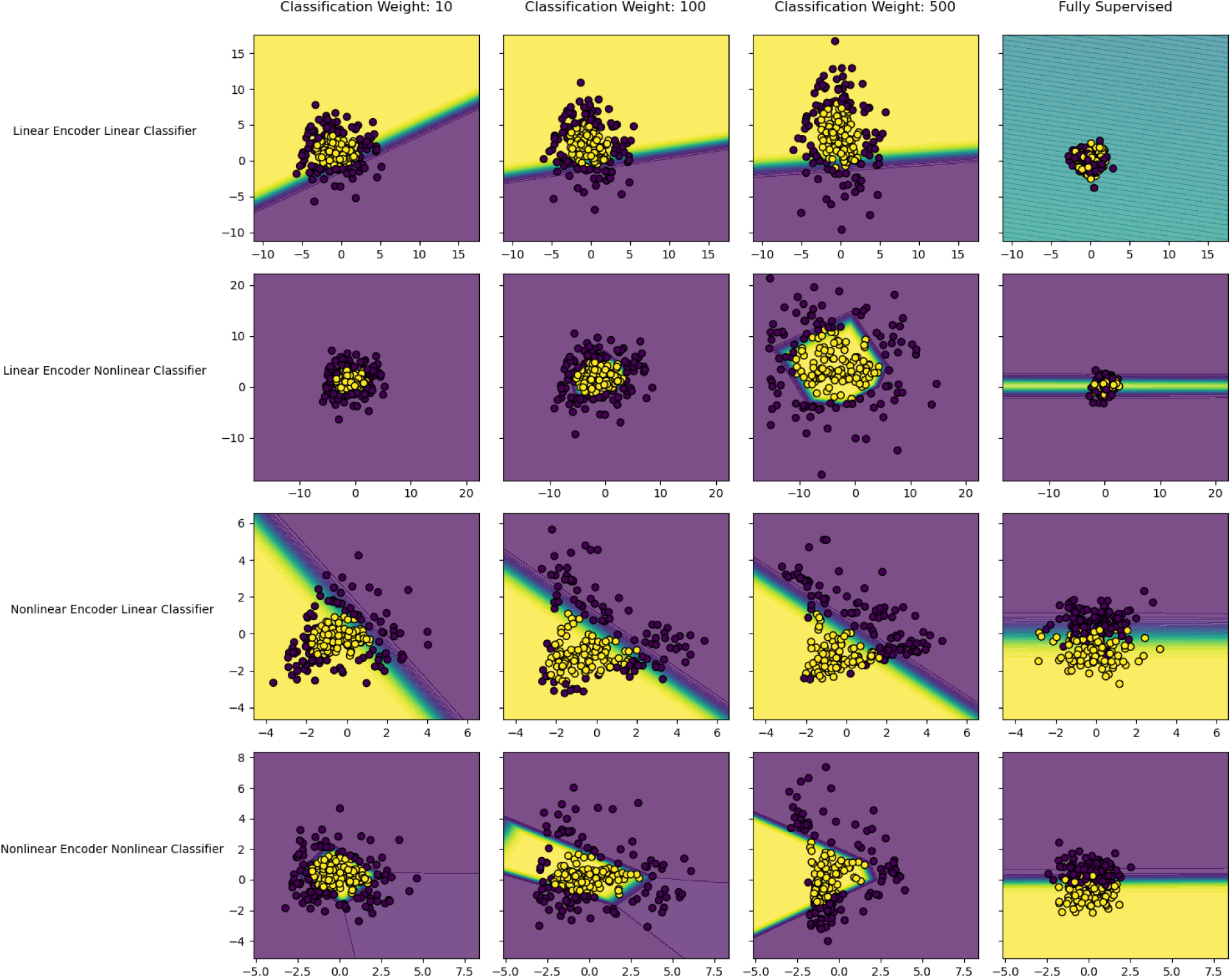
Learned Latent Spaces for Dataset 4. Each panel is the learned latent space for a different iteration of VAE-C model. Rows correspond to different architectures, while columns correspond to locations on the supervision continuum, with classification weights from left to right (10, 100, 500, and fully supervised). Each dot is the learned latent representation of a simulated fMRI observation. The color of the dot is the “observed” (simulated) behavioral label for that fMRI observation. The background color of each panel represents the models’ prediction of behavior labels for that location in the latent space. Model predictions are probabilistic so areas in which one class is not highly probable show graded coloration. For dataset 4, the latent spaces change between columns 3 and 4 as models transition to fully supervised models. Fully supervised models latent spaces fail to represent the many to one mappings present in the ground-truth simulated data. Next for the linear encoder, linear classifier (first row) the graded coloration and poor classification performance demonstrates the model’s inability to learn the nonlinear separation between behavioral labels. Finally, both models with nonlinear encoders the latent space becomes progressively distorted along the supervision continuum.

**Table 5.**
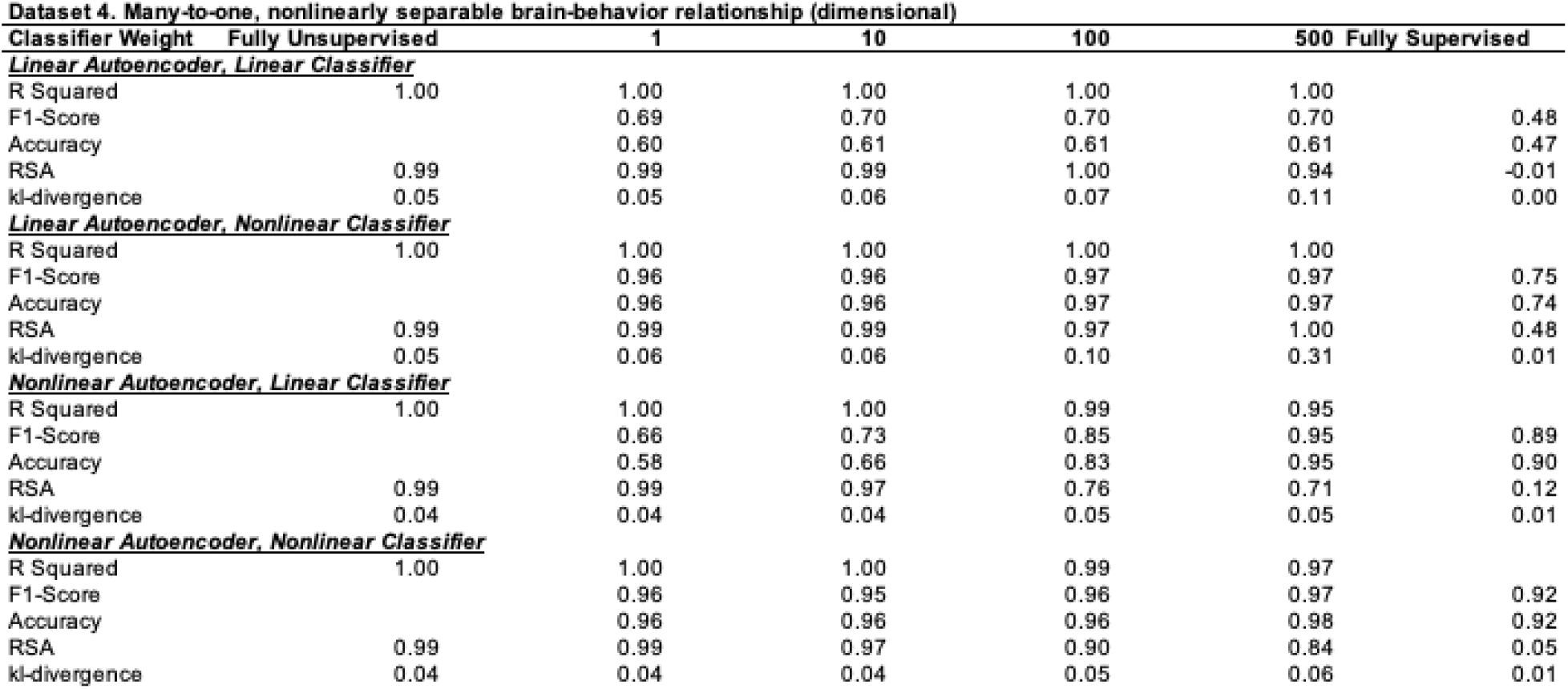
Columns of this table are the location on the unsupervised to the supervised continuum models were trained at. Rows are grouped by the VAE-C architecture a model was trained with. Within each architectural group, rows correspond to models’ performance in terms of reconstruction (R2), classification (F1-Score), and similarity of the learned and ground truth latent space (Representational Similarity Analysis; RSA). In dataset 4 all models continued to reconstruct well. Models with nonlinear classifiers generally classified well. All unsupervised and semi-supervised models performed well in terms of Representational Similarity, but fully supervised models performed extremely poorly. he nonlinear autoencoder, linear classifier model showed an interesting set of results, as the classification weight increased the F1-Score increased but representational similarity decreased, suggesting with increasing classification weight this model learned distorted latent spaces to accommodate the linear classifier. These changes were accompanied by a slight decrease in reconstruction accuracy (R2). The nonlinear/nonlinear model also showed a similar set of results where despite classifying well at all levels of supervision as the supervised weight increased the model showed slight decrements in representational similarity (RSA) and reconstruction performance (R2) suggesting distortions of the latent space.

**Table 6.** Summary of results across all datasets and framework variants. Modeling architectures (along rows) had either linearly or nonlinearly capable encoders and/or classifiers. The brain-behavior relationships in the simulated datasets (along columns) were linearly or nonlinearly separable and also had one-to-one or different versions of many-to-one mappings. Model architectures can “match” (green highlighted cells) or “violate” (red highlighted cells) the data generative process of the ground truth. “Allowable” (yellow highlighted cells) architectures are flexible enough to model the data even if they are not a direct match in terms of the linearity of the encoder and/or the classifier to the data generative process. Each cell summarizes the pattern of results in terms of reconstruction performance (R2), classification performance (F1-Score), and similarity of the structure of learned latent representations to the ground truth latent space (Pearson’s r) for a given model architecture and dataset, across unsupervised to supervised continuum. For the linearly separable one to one mapping dataset, all architectural variants performed consistently across all points in the supervision continuum, with high reconstruction (R2), classification (F1-Score), and just moderate decreases in the representational similarity to the ground truth in the fully supervised models (Pearson’s r). For the linearly separable, many to one mapping dataset models again had high reconstruction (R2) and classification (F1-Score) performance across all architectural variants for across all points in the supervision continuum. The representational similarity (Pearson’s r) however showed greater decreases for all architectures as the models were trained with greater supervision. For the non linearly separable, many to one, cluster-like mapping dataset all architectural variants had high reconstruction performance (R2), however only models with nonlinear classifiers had consistently high classification performance (F1-Score). The fully linear architecture had low classification performance (F1-Score) across all points in the supervision continuum, while the nonlinear autoencoder, linear classifier model had low classification performance with low degrees of supervision and high performance with high levels of supervision. Models with linear encoders had high representational similarity (Pearson’s r) for semi-supervised variants while models with nonlinear encoders had small decrements in representational similarity with increasing supervision. All architectures had low representational similarity when trained in a fully supervised way. The results for the nonlinearly separable, many to one, continuous, dataset mirrored the results described above for the nonlinearly separable many to one cluster like dataset.

|  |  | **Brain - Behavior Relationship** |  |  |  |
| --- | --- | --- | --- | --- | --- |
|  |  | **Linearly Separable, One to One** | **Linearly Separable, Many to One** | **Nonlinearly Separable, Many to One (Cluster-like)** | **Nonlinearly Separable, Many to One (Continuous)** |
| AutoEncoder Architecture | Classifier Architecture |  |  |  |  |
| Linear | Linear | **r2:** high<br>**F1-Score:** high<br>**Pearson's r:** moderate<br>decreases in fully supervised model | **r2:** high<br>**F1-Score:** high<br>**Pearson's r:** decreases as weight increases | **r2:** high<br>**F1-Score:** low<br>**Pearson's r:** Large decreases in fully supervised models | **r2:** high<br>**F1-Score:** low<br>**Pearson's r:** Large decreases in fully supervised models |
| Linear | Nonlinear | **r2:** high<br>**F1-Score:** high<br>**Pearson's r:** moderate<br>decrease in fully supervised model | **r2:** high<br>**F1:** high<br>**Pearson's r:** decreases as supervision increases | **r2:** high<br>**F1-Score:** high<br>**Pearson's r:** Large decreases in fully supervised models | **r2:** high<br>**F1-Score:** high<br>**Pearson's r:** Large decreases in fully supervised models |
| Nonlinear | Linear | **r2:** high<br>**F1-Score:** high<br>**Pearson's r:** moderate<br>decrease in fully supervised model | **r2:** high<br>**F1-Score:** high<br>**Pearson's r:** decreases as supervision increases | **r2:** high<br>**F1-Score:** increases from low to high with as supervision increases<br>**Pearson's r:** small decreases with increasing supervision, large decrease in fully supervised model | **r2:** high<br>**F1-Score:** increases from low to high with as supervision increases<br>**Pearson's r:** small decreases with increasing supervision, large decrease in fully supervised model |
| Nonlinear | Nonlinear | **r2:** high<br>**F1-Score:** high<br>**Pearson's r:** moderate<br>decrease in fully supervised model | **r2:** high<br>**F1-Score:** high<br>**Pearson's r:** decreases as supervision increases | **r2:** high<br>**F1-Score:** high<br>**Pearson's r:** small decreases with increasing supervision, large decrease in fully supervised model | **r2:** high<br>**F1-Score:** high<br>**Pearson's r:** small decreases with increasing supervision, large decrease in fully supervised model |

## Discussion

In cognitive neuroscience, it is important to model both the distribution of brain states and how these brain states relate to behavior. Conventional unsupervised and supervised analytical approaches tend to focus on one or the other objective. Each offers an incomplete account of brain-behavior relations that alone may lead to erroneous conclusions. Attempts at replication will not protect against this situation unless studies are explicitly designed and data are analyzed guided by alternative sets of assumptions. Even then, when supervised and unsupervised analyses are conducted separately and compared, it remains unclear how to best interpret inconsistent results (e.g., Azari et al., 2020). A fully supervised, purely discriminative model might classify behavior more accurately, but might not: A fully supervised, purely discriminative model would not have the advantage of using learned latent dimensions as features that could be more informative for classification (as offered by the VAE-C framework). Various unsupervised methods (such as clustering) could be dominated by variance in the BOLD signal data that is not necessarily relevant to behavior. A better solution is to systematically model data under varied assumptions at the outset, which requires a modeling framework that is explicitly designed for this purpose. Our VAE-C approach is one such framework. It combines two key components (1) a VAE-C neural network architecture that simultaneously models the distribution of brain states and their mappings to behavior and (2) a model comparison strategy that, akin to a multiverse analysis (Steegen et al., 2016), varies the degree of linearity in architecture and the amount of weight given to the supervised objective. This framework allows scientists to discover whether more complex relations might be present and to adjust their modeling approaches and inferences accordingly.

Most published research in cognitive neuroscience assumes that brain-behavior relationships are one-to-one and linearly separable. When we used the VAE-C framework with data simulated for this ground truth (dataset 1), all variants in our framework produced similar results. All variants within the 2 (linearly or nonlinearly capable auto-encoder) x 2 (a linearly or nonlinearly capable classifier) x W (a weight ratio, W, for the learning objectives from fully unsupervised to fully supervised) framework produced consistent R2, F1, and Pearson’s r indices, indicating that all variants showed strong agreement in their reconstruction of the data, their accuracy in predicting behavioral labels, and the structure of the latent space, respectively^1^. By implication, any analysis that produces a consistent set of results across all three outcomes suggests that an assumption of one-to-one linear relations between brain and behavior is warranted.

A growing body of evidence suggests, however, that brain-behavior relationships are complex and consist of nonlinear, many-to-one mappings (Doyle et al., 2022; Marder & Taylor, 2011; Mizusaki & O’Donnell, 2021; Price & Friston, 2002; Sajid et al., 2023; Wang et al., 2024). The VAE-C framework provides evidence of such nonlinearities when architectures with nonlinearities improve F1-scores, indicating improved classification accuracies, as exemplified in the results for data simulated for nonlinear mappings (datasets 3 and 4). The VAE-C framework also provides a powerful, flexible toolkit for identifying and analyzing many-to-one brain-behavior relationships for cases represented by the simulated datasets. Evidence for such many-to-one mappings occurs when large shifts in latent space organization are observed as models transition from unsupervised to supervised learning, as illustrated by changes in Pearson’s *r* for datasets 2, 3, and 4. When shifts are observed, the framework offers several additional ways to further interrogate the nature of many-to-one relations. For example, a clustering method could be applied to the latent representations associated with the behavioral labels. Datasets 2 and 3, which were simulated to represent a many-to-one cluster-like structure and a subsequent cluster analysis would produce multiple clusters per class. It is also possible to sample multiple points in the latent space and pass them through the decoding portion of the autoencoder network to reconstruct the associated set of BOLD measurements. This process produces an estimate of the BOLD signal correlates associated with each cluster and the degree of variance in those correlates.

This latter analysis is even useful to characterize the BOLD signal correlates and variation in those correlates when one-to-one mapping is observed. A one-to-one linear relation between brain and behavior is a mapping of one behavioral category to *a distribution of brain states*, not to a single brain state. To estimate the distribution of BOLD signal patterns that correspond to a given behavioral category, multiple points in the latent space for that category can be sampled and passed through the decoding portion of the autoencoder network. Each point corresponds to a trial on a given run for a given participant for the given category. This process, therefore, produces a distribution of reconstructed BOLD signal patterns on a trial-by-trial basis. The central tendency of this distribution of reconstructed patterns would correspond to the BOLD signal pattern expected from a univariate GLM (i.e., a standard modeling approach used in fMRI analyses). This single pattern is an abstraction, however, that need not be observed for *any* individual trial of *any* individual participant (see Clark-Polner et al., 2017; Westfall et al., 2017; Westlin et al., 2023). Other moments of the distribution, such as its variance, skew, and kurtosis are also important for characterizing the brain-behavior relationship. For example, one possibility is that there is little variance around the central tendency, making the central tendency a justified estimate of what the brain is doing during instances of that behavioral category. In such cases, it’s a reasonable approximation to infer that the behavioral category corresponds to a single brain state. This is the standard assumption made in most studies that assume a one-to-one linear brain-behavior mapping. It’s also possible that the variance is substantial and reproducible and therefore cannot be dismissed when considering the brain-behavior relationship. The autoencoder portion of the VAE-C framework learns only reliable variance and reconstructs the variance that the model can learn. In such cases, it’s not justified to infer that a behavioral category corresponds to a single brain state. (Model overfitting is minimized by both the information bottleneck and variational aspects of the architecture (ref) and can be detected by the difference between training and testing model statistics, such as the R2 scores). A one-to-one linear mapping with a wide (platykurtic) distribution vs. a narrow (leptokurtic) distribution of reconstructed BOLD signal patterns would obviously be interpreted very differently.

## Conclusion

The VAE-C framework is a versatile tool for evaluating assumptions about brain-behavior relationships during data modeling. Researchers are empowered to explicitly test their assumptions before drawing inferences from their findings. By integrating supervised and unsupervised objectives, the VAE-C framework offers insights that traditional methods obscure. Using four datasets that were simulated to represent four different ground-truth brain-behavior relationships, we tested two common assumptions – linearity and one-to-one mapping – that currently dominate much of cognitive neuroscience. Other assumptions, such as the time-invariant nature of brain-behavior mappings, should also be tested. Ultimately, frameworks like VAE-C will uncover the diversity and richness of brain-behavior relations. Only with such discoveries will neuroscience produce more robust, justified knowledge about brain-behavior relations.

## Acknowledgements

This paper was written with generous support from the National Science Foundation (BCS 1947972), the National Institutes of Health (R01 AG071173, R01 MH109464, R01 MH113234, R21 MH129902, R21 AG080198, RF1 AG078340, and R56 AG071486), the U.S. Army Research Institute for the Behavioral and Social Sciences (W911NF-16-1-019), and the Unlikely Collaborators Foundation. The views, opinions, and/or findings contained in this manuscript are those of the authors and shall not be construed as an official Department of the Army position, policy, or decision, unless so designated by other documents, nor do they necessarily reflect the views of the Unlikely Collaborators Foundation.

## Data and Code availability

All data and code for this manuscript can be found online at: https://github.com/kcmcveigh/vae-c_simulations

## Supplemental

### Additional Modeling Details

The KL term in the *L_VAE_* acts as a regularization term that tries to make sure that the posterior *P*(*z|x*) appear closer to prior *P*(*z*) i.e. *N*(0,*I*). The benefit of adding the KL term is that it introduces the completeness and continuity in the latent space hence allowing the notion of gradient over information embedded in the latent space *P*(*z*). This results in natural disentanglement of the information encoded in the latent space.

### Neural Network Hyperparameters

Our primary goal in comparing several neural network architectures is to show more expressive networks (i.e., nonlinear versus linear) allows for modeling a broader range of brain-behavior relationships. Since neural networks are universal function approximators, we do not consider our hyperparameter choices for the nonlinear models to confer any special properties of these networks, and believe a wide range of other architectures would perform similarly (i.e. with different number of hidden layers or number of neurons per layer). It is common in machine learning to explore a wide range of architectures to optimize performance. However, the architectures we report were the first we tested, and they performed near-optimally, so we did not pursue an exhaustive parameter search.

Our hyperparameter choices were as follows:

- For the nonlinear encoder, we used a single hidden layer with 25 neurons and a Tanh activation function. Given that our data were generated with a linear mapping from the latent space to fMRI representations, we chose this simple architecture. Tanh’s output range includes negative values, which aligns with the fMRI data’s range.
- For the nonlinear classifier, we used two hidden layers of sizes four and three, with ReLU activations. We added a second layer because research has shown that increasing the number of layers can improve the model’s ability to learn complex functions. While the relationships in our simulations are relatively simple, we believed the additional layer would help capture the structure. ReLU activations were chosen due to their proven success in classification tasks.

Additionally the set of classification weights were chosen to sample a range a of weights from no weight on the classification objective to all weight on the classification. To do this we used the following set of weights [ 0, 1, 10, 100, 500, Fully Supervised].

## Simulated Datasets

### Overview

In this study, we conducted four simulations to evaluate our model’s performance across various brain-behavior mapping conditions. The simulations generated synthetic data representing latent brain states, behavioral labels, and fMRI observations. The latent brain states were modeled as points in a 2-dimensional space, sampled from Gaussian distributions. Behavioral labels were generated based on the class membership of these points, and fMRI data was generated by projecting the latent points into a higher-dimensional space using a randomly initialized linear transformation. The parameters used for each simulation are outlined below.

### Dataset 1: Linearly Separable Gaussian Classes

When simulating dataset 1, latent brain states were sampled from two distinct Gaussian distributions in a 2-dimensional space. A total of 1000 data points were generated, with 500 points sampled from each Gaussian. The behavioral labels were assigned such that points from the first Gaussian were labeled as “Class A” and points from the second Gaussian as “Class B.” These latent points were then projected into a 100-dimensional fMRI space using a randomly generated projection matrix.

### Dataset 2: Linearly Separable Many-to-One Mapping

When simulating dataset 2, latent brain states were sampled from four distinct Gaussian distributions, representing a more complex mapping scenario. A total of 1000 data points were sampled, with 250 points drawn from each of the four distributions. Behavioral labels were assigned such that data from two of the distributions were mapped to “Class A” and the other two distributions were mapped to “Class B.” The latent points were again projected into a 100-dimensional fMRI space using the same randomly generated projection matrix as in Simulation 1.

### Dataset 3: Non-Linearly Separable Many-to-One Mapping

When simulating dataset 3, we modeled a nonlinearly separable many-to-one brain-behavior relationship. Latent brain states were again sampled from four distinct Gaussian distributions, with 250 points sampled from each distribution. However, in this simulation, the class labels were assigned such that the data points could not be separated by a simple linear decision boundary. The fMRI data were generated by projecting the latent points into 100-dimensional space using the same projection matrix as before.

Dataset 4: Non-Linearly Separable Many-to-One Mapping

When simulating dataset 4, latent brain states were sampled from a single Gaussian distribution in the 2-dimensional space. The behavioral labels were generated by assigning the 50% of data points closest to the mean of the Gaussian to “Class A” and the remaining points to “Class B,” following the “Anna Karenina” model. The fMRI data were again generated by projecting the latent data points into a 100-dimensional space using the same projection matrix as in the previous simulations.

### Projection Matrix for fMRI Data Generation

In all simulations, the latent points were projected into 100-dimensional fMRI space using a random projection matrix, this projection matrix is available on the project github

---

**Table 1:** Parameters for Dataset 1.

| Parameter | Class A (Gaussian 1) | Class B (Gaussian 2) |
| --- | --- | --- |
| Mean | [0.0763, 0.7799] | [2.0763, 0.7799] |
| Covariance Matrix | [[0.4384, 0], [0, 0.7235]] | [[0.4384, 0], [0, 0.7235]] |

**Table 2:** Parameters for Dataset 2.

| Parameter | Class A (Dist 1) | Class A (Dist 2) | Class B (Dist 1) | Class B (Dist 2) |
| --- | --- | --- | --- | --- |
| Mean | [0.7182, 0.1664] | [6.7182, 0.1664] | [0.7182, 3.1664] | [6.7182, 3.1664] |
| Covariance Matrix | [[1.4280, 0], [0, 0.6255]] | [[1.1753, 0], [0, 0.5266]] | [[1.4280, 0], [0, 0.6255]] | [[1.1753, 0], [0, 0.5266]] |

**Table 3:**
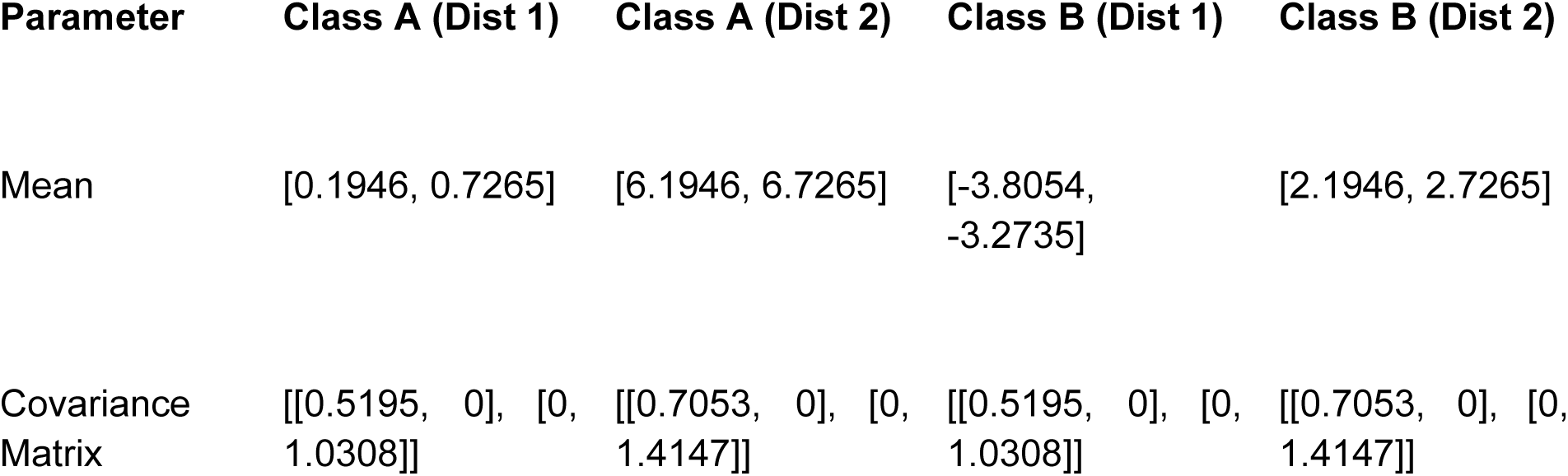
Parameters for Dataset 3.

| Parameter | Class A (Dist 1) | Class A (Dist 2) | Class B (Dist 1) | Class B (Dist 2) |
| --- | --- | --- | --- | --- |
| Mean | [0.1946, 0.7265] | [6.1946, 6.7265] | [-3.8054, -3.2735] | [2.1946, 2.7265] |
| Covariance Matrix | [[0.5195, 0], [0, 1.0308]] | [[0.7053, 0], [0, 1.4147]] | [[0.5195, 0], [0, 1.0308]] | [[0.7053, 0], [0, 1.4147]] |

**Table 4:** Parameters for Dataset 4.

| Parameter | Value |
| --- | --- |
| Mean | [0.9482, 0.0547] |
| Covariance Matrix | [[1.1911, 0], [0, 0.8295]] |

## Footnotes

1 Note that in our simulations Pearson’s r was used to quantify the relationship between the ground truth latent structure, and the modeled latent structure. This comparison is not possible in empirical datasets, however relative changes in the structure of the latent space can be compared by using the unsupervised model as an anchor against which changes in the latent space can be measured.

## References

Azari, B., Westlin, C., Satpute, A. B., Hutchinson, J. B., Kragel, P. A., Hoemann, K., Khan, Z., Wormwood, J. B., Quigley, K. S., Erdogmus, D., Dy, J., Brooks, D. H., & Barrett, L. F. (2020). Comparing supervised and unsupervised approaches to emotion categorization in the human brain, body, and subjective experience. Scientific Reports, 10(1), 20284. 10.1038/s41598-020-77117-8

Barrett, L. F. (2022). Context reconsidered: Complex signal ensembles, relational meaning, and population thinking in psychological science. The American Psychologist, 77(8), 894–920. 10.1037/amp0001054

Barrett, L. F., & Satpute, A. B. (2013). Large-scale brain networks in affective and social neuroscience: Towards an integrative functional architecture of the brain. Current Opinion in Neurobiology, 23(3), 361–372. 10.1016/j.conb.2012.12.012

Bethge, D., Hallgarten, P., Grosse-Puppendahl, T., Kari, M., Chuang, L. L., Özdenizci, O., & Schmidt, A. (2022). EEG2Vec: Learning Affective EEG Representations via Variational Autoencoders. 2022 IEEE International Conference on Systems, Man, and Cybernetics (SMC), 3150–3157. 10.1109/SMC53654.2022.9945517

Bishop, C. M. (2006). Pattern recognition and machine learning. Springer

Buzsáki, G. (2019). The Brain from Inside Out. Oxford University Press.

Bzdok, D., Eickenberg, M., Grisel, O., Thirion, B., & Varoquaux, G. (2015). Semi-Supervised Factored Logistic Regression for High-Dimensional Neuroimaging Data. Advances in Neural Information Processing Systems, 28. https://proceedings.neurips.cc/paper/2015/hash/06a15eb1c3836723b53e4abca8d9b879-Abstract.html

Clark-Polner, E., Johnson, T. D., & Barrett, L. F. (2017). Multivoxel Pattern Analysis Does Not Provide Evidence to Support the Existence of Basic Emotions. Cerebral Cortex, 27(3), 1944–1948. 10.1093/cercor/bhw028

Doyle, C. M., Lane, S. T., Brooks, J. A., Wilkins, R. W., Gates, K. M., & Lindquist, K. A. (2022). Unsupervised classification reveals consistency and degeneracy in neural network patterns of emotion. Social Cognitive and Affective Neuroscience, 17(11), 995–1006. 10.1093/scan/nsac028

Dubova, M., & Goldstone, R. L. (2023). Carving joints into nature: Reengineering scientific concepts in light of concept-laden evidence. Trends in Cognitive Sciences, 27(7), 656–670. 10.1016/j.tics.2023.04.006

Edelman, G. M., & Gally, J. A. (2001). Degeneracy and complexity in biological systems. Proceedings of the National Academy of Sciences, 98(24), 13763–13768. 10.1073/pnas.231499798

Ezekiel, M. (1930). Methods of correlation analysis (pp. xiv, 427). Wiley.

Finn, E. S., Glerean, E., Khojandi, A. Y., Nielson, D., Molfese, P. J., Handwerker, D. A., & Bandettini, P. A. (2020). Idiosynchrony: From shared responses to individual differences during naturalistic neuroimaging. NeuroImage, 215, 116828. 10.1016/j.neuroimage.2020.116828

Fukushima, K. (1980). Neocognitron: A self-organizing neural network model for a mechanism of pattern recognition unaffected by shift in position. Biological Cybernetics, 36(4), 193–202. 10.1007/BF00344251

Goutte, C., Toft, P., Rostrup, E., Nielsen, F. Å., & Hansen, L. K. (1999). On Clustering fMRI Time Series. NeuroImage, 9(3), 298–310. 10.1006/nimg.1998.0391

Han, K., Wen, H., Shi, J., Lu, K.-H., Zhang, Y., Fu, D., & Liu, Z. (2019). Variational autoencoder: An unsupervised model for encoding and decoding fMRI activity in visual cortex. NeuroImage, 198, 125–136. 10.1016/j.neuroimage.2019.05.039

Hastie, T., Friedman, J., & Tibshirani, R. (2001). Unsupervised Learning. In T. Hastie, J. Friedman, & R. Tibshirani (Eds.), The Elements of Statistical Learning: Data Mining, Inference, and Prediction (pp. 437–508). Springer. 10.1007/978-0-387-21606-5_14

Haxby, J. V. (2012). Multivariate pattern analysis of fMRI: The early beginnings. NeuroImage, 62(2), 852–855. 10.1016/j.neuroimage.2012.03.016

Haynes, J.-D., & Rees, G. (2006). Decoding mental states from brain activity in humans. Nature Reviews Neuroscience, 7(7), 523–534. 10.1038/nrn1931

Hornik, K., Stinchcombe, M., & White, H. (1989). Multilayer feedforward networks are universal approximators. Neural Networks, 2(5), 359–366. 10.1016/0893-6080(89)90020-8

Huang, H., Hu, X., Zhao, Y., Makkie, M., Dong, Q., Zhao, S., Guo, L., & Liu, T. (2018). Modeling Task fMRI Data Via Deep Convolutional Autoencoder. IEEE Transactions on Medical Imaging, 37(7), 1551–1561. IEEE Transactions on Medical Imaging. 10.1109/TMI.2017.2715285

Khan, Z., Wang, Y., Sennesh, E., Dy, J., Ostadabbas, S., van de Meent, J.-W., Hutchinson, J. B., & Satpute, A. B. (2022). A computational neural model for mapping degenerate neural architectures. Neuroinformatics, 1–15.

Kingma, D. P., Rezende, D. J., Mohamed, S., & Welling, M. (2014). *Semi-Supervised Learning with Deep Generative Models* (arXiv:1406.5298). arXiv. 10.48550/arXiv.1406.5298

Kragel, P. A., Koban, L., Barrett, L. F., & Wager, T. D. (2018). Representation, Pattern Information, and Brain Signatures: From Neurons to Neuroimaging. Neuron, 99(2), 257–273. 10.1016/j.neuron.2018.06.009

Kriegeskorte, N., Mur, M., & Bandettini, P. A. (2008). Representational similarity analysis—Connecting the branches of systems neuroscience. Frontiers in Systems Neuroscience, 2. 10.3389/neuro.06.004.2008

Lee, K. M., Ferreira-Santos, F., & Satpute, A. B. (2021). Predictive processing models and affective neuroscience. Neuroscience & Biobehavioral Reviews, 131, 211–228. 10.1016/j.neubiorev.2021.09.009

Lindquist, K. A., Jackson, J. C., Leshin, J., Satpute, A. B., & Gendron, M. (2022). The cultural evolution of emotion. Nature Reviews Psychology, 1(11), 11. 10.1038/s44159-022-00105-4

Marder, E., & Taylor, A. L. (2011). Multiple models to capture the variability in biological neurons and networks. Nature Neuroscience, 14(2), 133–138. 10.1038/nn.2735

Mitchell, T. M., Hutchinson, R., Niculescu, R. S., Pereira, F., Wang, X., Just, M., & Newman, S. (2004). Learning to Decode Cognitive States from Brain Images. Machine Learning, 57(1), 145–175. 10.1023/B:MACH.0000035475.85309.1b

Mizusaki, B. E. P., & O’Donnell, C. (2021). Neural circuit function redundancy in brain disorders. Current Opinion in Neurobiology, 70, 74–80. 10.1016/j.conb.2021.07.008

Nakuci, J., Yeon, J., Kim, J.-H., Kim, S.-P., & Rahnev, D. (2023). Multiple brain activation patterns for the same task. BioRxiv. https://www.ncbi.nlm.nih.gov/pmc/articles/PMC10104176/

Norman, K. A., Polyn, S. M., Detre, G. J., & Haxby, J. V. (2006). Beyond mind-reading: Multi-voxel pattern analysis of fMRI data. Trends in Cognitive Sciences, 10(9), 424–430. 10.1016/j.tics.2006.07.005

O’Toole, A. J., Jiang, F., Abdi, H., Pénard, N., Dunlop, J. P., & Parent, M. A. (2007). Theoretical, Statistical, and Practical Perspectives on Pattern-based Classification Approaches to the Analysis of Functional Neuroimaging Data. Journal of Cognitive Neuroscience, 19(11), 1735–1752. 10.1162/jocn.2007.19.11.1735

Peelen, M. V., & Downing, P. E. (2023). Testing cognitive theories with multivariate pattern analysis of neuroimaging data. Nature Human Behaviour, 7(9), 1430–1441. 10.1038/s41562-023-01680-z

Pereira, F., Mitchell, T., & Botvinick, M. (2009). Machine learning classifiers and fMRI: A tutorial overview. NeuroImage, *45*(1, Supplement 1), S199–S209. 10.1016/j.neuroimage.2008.11.007

Poldrack, R. A. (2008). The role of fMRI in Cognitive Neuroscience: Where do we stand? Current Opinion in Neurobiology, 18(2), 223–227. 10.1016/j.conb.2008.07.006

Poldrack, R. A., Halchenko, Y. O., & Hanson, S. J. (2009). Decoding the Large-Scale Structure of Brain Function by Classifying Mental States Across Individuals. Psychological Science, 20(11), 1364–1372. 10.1111/j.1467-9280.2009.02460.x

Price, C. J., & Friston, K. J. (2002). Degeneracy and cognitive anatomy. Trends in Cognitive Sciences, 6(10), 416–421.

Putnam, H. (1960). Minds and Machines. In S. Hook (Ed.), Dimensions Of Mind: A Symposium. (pp. 138–164). NEW YORK University Press.

Rijsbergen, C. J. V. (1979). Information Retrieval (2nd ed.). Butterworth-Heinemann.

Rugg, M. D., & Thompson-Schill, S. L. (2013). Moving Forward With fMRI Data. Perspectives on Psychological Science, 8(1), 84–87. 10.1177/1745691612469030

Sajid, N., Gajardo-Vidal, A., Ekert, J. O., Lorca-Puls, D. L., Hope, T. M. H., Green, D. W., Friston, K. J., & Price, C. J. (2023). Degeneracy in the neurological model of auditory speech repetition. Communications Biology, 6(1), 1–10. 10.1038/s42003-023-05515-5

Sennesh, E., Khan, Z., Wang, Y., Hutchinson, J. B., Satpute, A., Dy, J., & van de Meent, J.-W. (2020). Neural Topographic Factor Analysis for fMRI Data. In H. Larochelle, M. Ranzato, R. Hadsell, M. F. Balcan, & H. Lin (Eds.), Advances in Neural Information Processing Systems (Vol. 33, pp. 12046–12056). Curran Associates, Inc. https://proceedings.neurips.cc/paper/2020/file/8c3c27ac7d298331a1bdfd0a5e8703d3-Paper.pdf

Steegen, S., Tuerlinckx, F., Gelman, A., & Vanpaemel, W. (2016). Increasing Transparency Through a Multiverse Analysis. Perspectives on Psychological Science, 11(5), 702–712. 10.1177/1745691616658637

Tong, F., & Pratte, M. S. (2012). Decoding Patterns of Human Brain Activity. Annual Review of Psychology, 63(Volume 63, 2012), 483–509. 10.1146/annurev-psych-120710-100412

Uttal, W. (2003). The New Phrenology. MIT Press.

Wang, Y., Kragel, P., & Satpute, A. B. (2024). Neural predictors of fear depend on the situation. Journal of Neuroscience. 10.1523/JNEUROSCI.0142-23.2024

Weaverdyck, M. E., Lieberman, M. D., & Parkinson, C. (2020). Tools of the Trade Multivoxel pattern analysis in fMRI: A practical introduction for social and affective neuroscientists. Social Cognitive and Affective Neuroscience, 15(4), 487–509. 10.1093/scan/nsaa057

Westfall, J., Nichols, T. E., & Yarkoni, T. (2017). Fixing the stimulus-as-fixed-effect fallacy in task fMRI. Wellcome Open Research, 1, 23. 10.12688/wellcomeopenres.10298.2

Westlin, C., Theriault, J. E., Katsumi, Y., Nieto-Castanon, A., Kucyi, A., Ruf, S. F., Brown, S. M., Pavel, M., Erdogmus, D., Brooks, D. H., Quigley, K. S., Whitfield-Gabrieli, S., & Barrett, L. F. (2023). Improving the study of brain-behavior relationships by revisiting basic assumptions. Trends in Cognitive Sciences, 27(3), 246–257. 10.1016/j.tics.2022.12.015

Zabihi, M., Kia, S. M., Wolfers, T., Boer, S. de, Fraza, C., Dinga, R., Arenas, A. L., Bzdok, D., Beckmann, C. F., & Marquand, A. (2024). Nonlinear latent representations of high-dimensional task-fMRI data: Unveiling cognitive and behavioral insights in heterogeneous spatial maps. PLOS ONE, 19(8), e0308329. 10.1371/journal.pone.0308329

